# DNT: Diploid Genomic Foundation Model

**DOI:** 10.64898/2026.09.05.749576

**Authors:** Guy Leib, Tal Zinger, Dan Ofer, Raizy Kellerman, Omri Nayshool, Dan Dominissini, Ariel Larey, Jeremy Levy, Yury Nahshan, Elay Dahan, Amit Bleiweiss, Nicole Bussola, Simon Lee, Shane O’Connell, Dung Hoang, Marissa Wirth, Noam D. Beckmann, Alexander W. Charney, Yoli Shavit, Nati Daniel, Gideon Rechavi

**Author notes:** These authors contributed equally (Guy Leib, Tal Zinger, Dan Ofer, Raizy Kellerman, Omri Nayshool).

## Abstract

Clinical interpretation of genetic variation depends on the diploid genotype, including zygosity, allele dosage and whether multiple variants occur in *cis* on the same homologue or in *trans* on different homologues. Most genomic language models process haploid sequences or combine independently encoded haplotypes downstream, so they do not directly represent the paired genotype in a single sequence. We introduce a reference-aligned diploid encoding for single-nucleotide variants (SNVs) and short insertions and deletions (indels), together with unphased and phase-retaining tokenizers that accept phased genotypes and convert them to single-sequence diploid representation. Using Nucleotide Transformer v3 backbones, we continue training 8-million- and 100-million-parameter models and evaluate an auxiliary Contrastive Phase Loss (CPL) designed to retain the phasing information of the variants in contextual representations. We evaluate on a novel compound-heterozygous benchmark containing 9,460 examples. Models whose inputs did not distinguish relative phase remained near chance, whereas our diploidic models improved discrimination with AUROC 0.649 ± 0.005, compared to 0.506 for the vocabulary-adapted control. These findings establish a method for making diploid genotype information accessible to genomic language models, rather than a universal improvement in variant prediction; validation in naturally observed, accurately phased clinical cohorts remains necessary.

## 1 Introduction

The human genome is diploid: each individual inherits one complete haploid complement from each parent, carried on homologous chromosomes. This duplication is not simple redundancy, because a variant can produce different phenotypes depending on whether it is present on one copy of a gene (heterozygous) or both (homozygous). In recessive diseases, the heterozygous person will mostly be considered a carrier rather than affected. In dominant diseases, even a single copy of the gene affected by a variant would be sufficient to demonstrate noticeable phenotype. When two variants occur on the same homologue they are in *cis*; when one occurs on each homologue they are in *trans*. Having this additional layer of complexity (compound heterozygous) is an example of the need to consider the phasing of detected variants, when investigating causes for genetic diseases[1–3] . These relationships are not absolute, as penetrance, variable expressivity, hypomorphic alleles, genetic modifiers and gene-specific mechanisms can alter the phenotypic consequences of a genotype[4].

Genomic foundation models (GFMs) have advanced through larger and more diverse pretraining corpora[5–7], long-context and state-space architectures[8–12], evolutionary supervision[13, 14], and sequence-to-function learning[10, 15, 16]. Most sequence-based models nevertheless operate on a haploid reference or an allele-substituted sequence. Variant effects are commonly estimated by inserting an alternative allele into the reference sequence and comparing the resulting prediction with that of the reference allele. This can characterize an individual genetic variant, but it cannot distinguish heterozygous from homozygous genotypes or determine whether two variants occur in *cis* or in *trans*.

Existing approaches capture complementary subsets of diploid information. Variant-panel and genotype-imputation models can represent phased haplotypes and learn linkage-disequilibrium structure, but generally operate over predefined marker sets rather than continuous nucleotide sequence[17–20]. Variation-aware sequence models retain local nucleotide context but typically describe population-level polymorphism rather than the two alleles carried by a particular individual[21, 22]. Reverse-complement-equivariant architectures process paired sequence orientations[23], but these correspond to the complementary DNA strands rather than the two inherited homologues. Recent zygosity-aware sequence models encode the two homologues separately and combine their representations downstream[24, 25]. Moreover, existing approaches do not jointly preserve explicit inserted and deleted nucleotide sequences together with diploid allelic state and haplotype orientation.

The methodological question is therefore not whether one representation of diploidy is intrinsically correct, but whether a sequence-native encoding that links the two alleles at each reference-aligned position provides a useful inductive bias for genotype-dependent prediction. The unresolved gap is a continuous nucleotide representation that jointly exposes the realized diploid state, preserves the sequence of short indels and, when phased genotypes are available, retains the assignment of alleles to homologues. A detailed model-by-model comparison is provided in Supplementary Note 1.

We address this gap with a reference-aligned diploid encoding designed for SNVs and short indels (Fig. 1a,b). Homozygous positions retain the standard nucleotide alphabet, whereas heterozygous positions are represented by atomic allele-pair symbols. Dedicated structural tokens mark SNVs and indel boundaries, and local alignment with allele-specific gap padding preserves inserted and deleted sequence base by base (Fig. 2). The resulting genotype is represented in one sequence stream and can be processed jointly by a single encoder.

**Figure 1.**
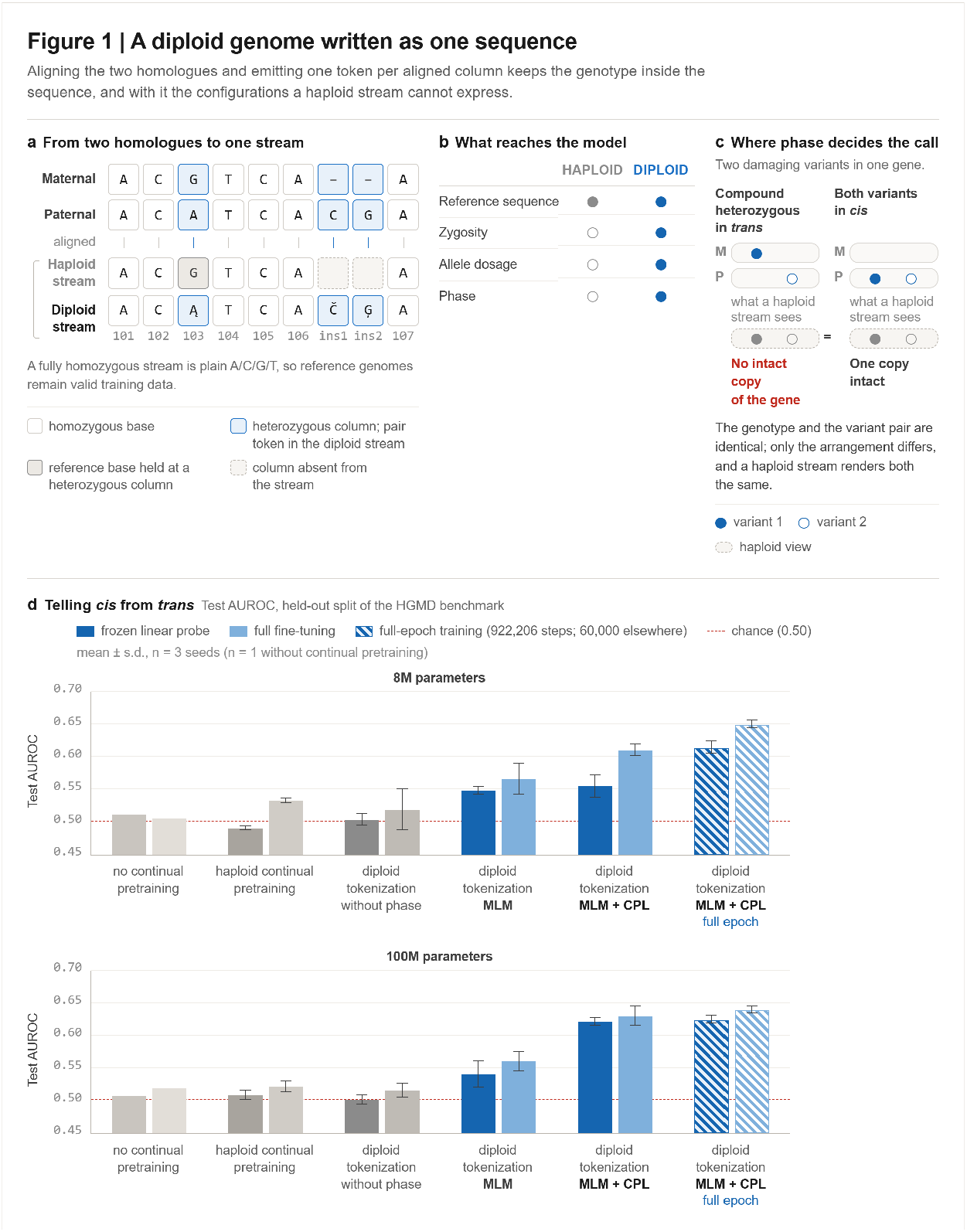
A diploid genome written as one sequence. Aligning the two homologues and emitting one token per aligned column keeps the genotype inside the sequence, together with the configurations a haploid stream cannot express. a. Reference-aligned diploid encoding. Maternal and paternal homologues are aligned column by column and emitted as a single token stream, one token per aligned position. Homozygous columns retain the standard nucleotide alphabet, whereas heterozygous columns are replaced by an atomic allele-pair symbol, so both alleles reach the encoder in a single forward pass: Ą encodes the G/A substitution at position 103, and Č and Ģ encode the two inserted columns. Allele-specific gap padding preserves inserted bases as sequence rather than collapsing them to a generic event token. A haploid stream instead holds the reference base at heterozygous columns and omits columns absent from the reference, admitting an alternate allele only by substitution, one variant per pass, so the two copies never appear together. A fully homozygous stream reduces to plain A/C/G/T, so reference genomes remain valid training data. b. Genotype properties available to the model under each encoding. A haploid stream conveys the reference sequence alone; the diploid stream additionally carries zygosity, allele dosage and, where phased input is supplied, the assignment of alleles to homologues. c. The configuration that phase determines. Two damaging variants in the same gene disrupt both copies when they lie in trans, whereas the same two variants in cis leave one copy intact. Genotype and variant pair are identical in the two cases and only the arrangement differs, so a haploid stream renders them indistinguishable. d. Compound-heterozygous cis versus trans classification across pretraining conditions, at 8 million (top) and 100 million (bottom) parameters. Bars show test AUROC on the held-out 1,811-example split of the HGMD-derived benchmark; dark bars are frozen linear probes, light bars full fine-tuning, and hatched bars the full-epoch schedule of 922,206 optimizer steps, against 60,000 for all other conditions. Error bars are s.d. over three probe seeds; the no-continual-pretraining control is a single run. The dashed line marks chance at 0.50. The vertical axis is truncated at 0.45, so bar length is not proportional to AUROC.

**Figure 2.**
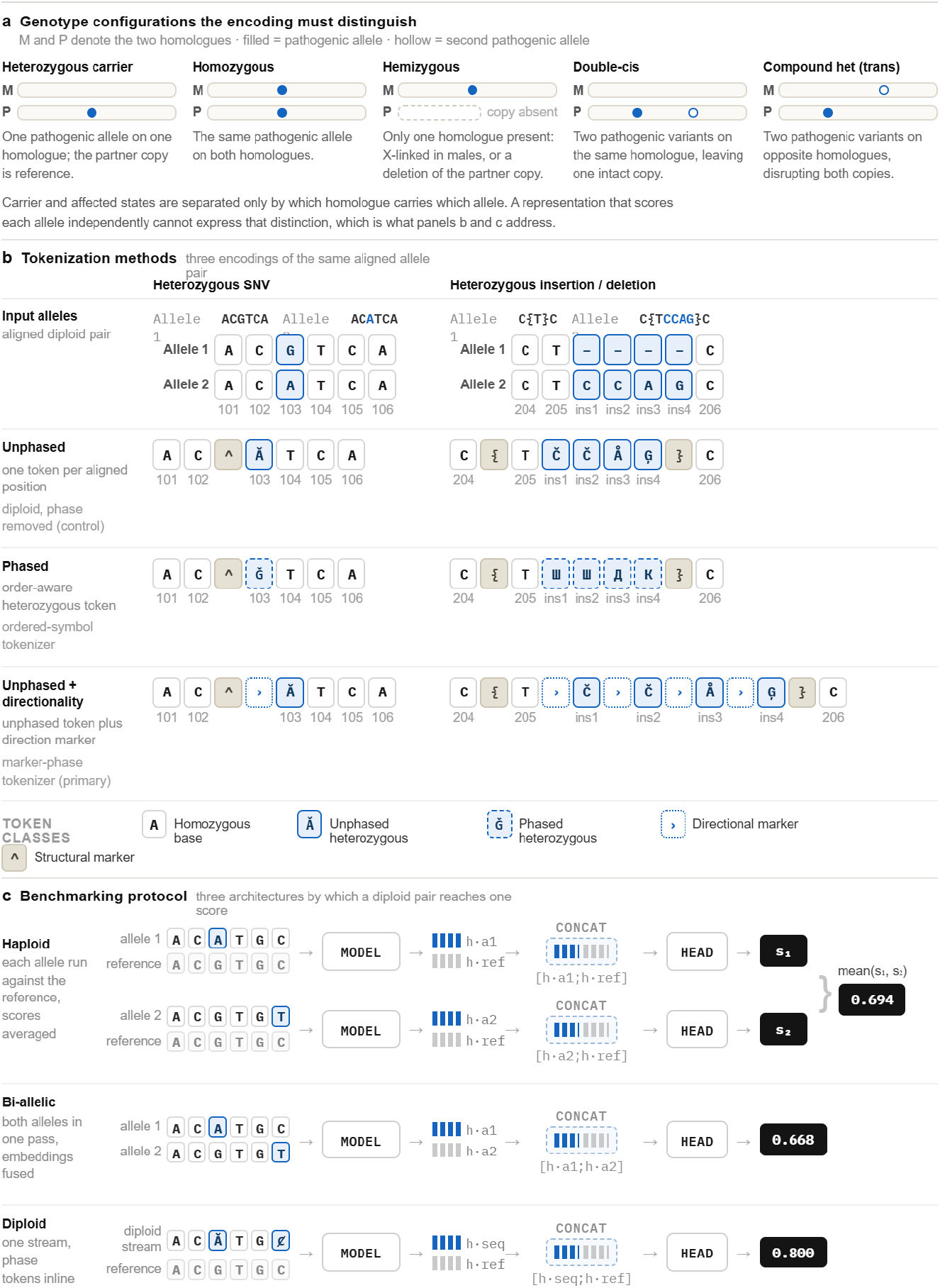
Representing diploid genotypes: tokenization and evaluation. One token per aligned column preserves the configurations a per-allele representation cannot express, and the readout architecture determines what a model can use. **a.** Genotype configurations the encoding must distinguish. M and P denote the two homologues; filled and hollow marks are the two pathogenic alleles. Heterozygous carrier, homozygous, hemizygous, double-*cis* and compound heterozygous (*trans*) states are separated only by which homologue carries which allele, and a representation that scores each allele independently cannot express that distinction. **b.** Three encodings of the same aligned allele pair, shown for a heterozygous single-nucleotide variant at position 103 (G / A) and for a four-base insertion carried by allele 2 within an indel span delimited by { and }. All three emit one token per aligned position, with the variant marker ^ preceding the heterozygous token. The unphased token Ⱥ is order-invariant, the phased token Ğ records allele order, and the marker-phase encoding prefixes the unphased token with an orientation marker. Gap-containing pairs become gap tokens, and phased symbols additionally record which allele carries the inserted sequence. Position labels beneath each token give the reference coordinate the token was emitted from; markers carry no coordinate. Token classes are distinguished by outline style as well as colour, so the figure reads in greyscale. **c.** The three readout architectures by which a diploid pair reaches a single score: the haploid readout runs each allele against the reference and averages the two scores, the bi-allelic readout passes both alleles in one forward pass and fuses their embeddings, and the diploid readout encodes the genotype as one stream with phase tokens inline. The complete token vocabulary is given in Supplementary Table 1.

We define three diploid tokenizers. The **unphased tokenizer** maps reciprocal allele pairs to the same symbol. The **ordered-phase tokenizer** assigns different symbols to the two allele orders. The **marker-phase tokenizer** uses an unordered allele-pair symbol together with an explicit orientation marker (Fig. 2). The phase-retaining tokenizers do not infer phase: they require phased input and preserve the haplotype assignments supplied by an upstream phasing procedure and conveyed in the VCF file. In this study, those assignments were obtained from statistically phased 1000 Genomes data[26, 27, 31] and may inherit phasing errors.

Because phase-retaining input does not guarantee that contextual representations will retain the supplied ordering, we additionally evaluate Contrastive Phase Loss (CPL), an auxiliary objective applied alongside masked language modeling. CPL is introduced as a representation-learning intervention rather than as a phasing method; its effect is tested by asking whether downstream models can recover the relative phase of variant pairs. The distinction between global homologue labelling and biologically meaningful relative phase is examined explicitly in the Results and Methods sections.

Using 8-million- and 100-million-parameter Nucleotide Transformer v3[28] backbones, we continually pretrain diploid models on phased population-scale human genomes and compare them with a haploid continual pretraining control, an unphased diploid model and a two-stream haplotype baseline (Fig. 1c). The primary evaluation is a compound-heterozygous benchmark in which *cis* and *trans* examples are drawn from the same recessive genes and the same individuals, and differ in whether the two injected variants lie on one homologue or on both (Fig. 1d). We complement this analysis with simplified mode-of-inheritance-aware ClinVar SNV and indel benchmarks and general haploid controls (Fig. 3).

**Figure 3.**
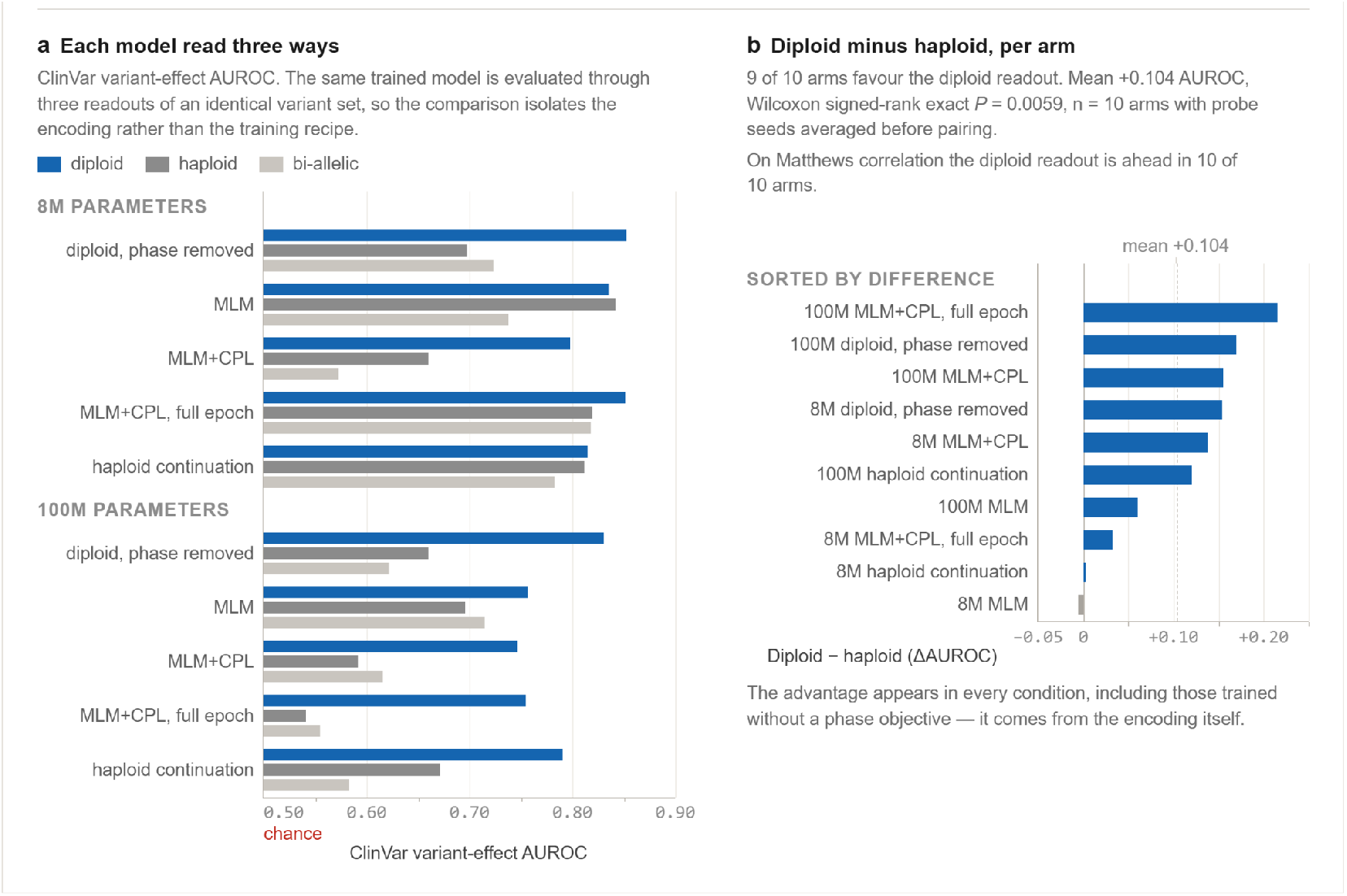
Diploid encoding improves clinical variant-effect prediction. Reading the same trained model through the diploid encoding raises ClinVar variant-effect prediction in almost every condition tested, including conditions trained without any phase objective. a. ClinVar supervised variant-effect AUROC at 8 million and 100 million parameters. Each trained model is read out three ways on an identical variant set: the diploid tokenization, a single haplotype, and both haplotypes supplied as two separate sequences. Because the variant set and the trained model are held fixed and only the readout changes, the comparison isolates the input encoding rather than the pretraining recipe. Conditions are the haploid continual-pretraining control, the phase-removed diploid model, and marker-phase models trained with masked language modelling alone, with Contrastive Phase Loss, and with Contrastive Phase Loss to a full epoch. Bars are the mean over three probe seeds and chance is 0.50. b. Paired difference between the diploid and haploid readouts for each arm, sorted by effect size. The diploid readout leads in 9 of 10 arms by a mean of 0.104 AUROC (Wilcoxon signed-rank exact P = 0.0059, n = 10 arms, probe seeds averaged within an arm before pairing). On Matthews correlation the diploid readout leads in 10 of 10 arms. The single exception on AUROC is the 8 million parameter marker-phase model trained with masked language modelling alone, where the haploid readout leads by 0.006. Because the advantage also appears in conditions trained without any phase objective, it follows from the encoding rather than from phase supervision.

Our primary contribution is the encoding scheme itself; we additionally release the pretrained model checkpoints and the benchmark suite and evaluation framework used to validate it. The present study tests whether genotype configuration remains accessible after contextual encoding; it does not claim clinical-grade variant interpretation or universal superiority over separate-haplotype approaches.

## 2 Results

### 2.1 Study design and reference aligned diploid encoding

The study separates three questions: whether both homologues can be represented within one continuous nucleotide stream, whether phase-retaining input survives continual pretraining, and whether the resulting representations support genotype-dependent prediction. We initialized 8-million- and 100-million-parameter NTv3 checkpoints, adapted their vocabularies, continually pretrained all model conditions on phased 1000 Genomes sequences and evaluated frozen and fine-tuned representations.

The reference-aligned diploid encoding represents the allele pair at each locus as one state (Fig. 2). The unphased tokenizer discards allele order, whereas the ordered-phase and marker-phase tokenizers preserve the supplied assignment of alleles to the two homologues. These names are used consistently throughout the manuscript. The phase-retaining tokenizers consume phased genotypes but do not estimate phase.

For short indels, the two realized alleles are aligned within explicit span boundaries and the shorter allele is padded with gap states (Fig. 2b). This preserves inserted or deleted nucleotides and their local coordinate relationship rather than reducing every indel to one generic event token. The encoding is currently limited to SNVs and short indels and does not represent structural variants or copy-number changes.

The union vocabulary defines 49 sequence and metadata symbols. Five nucleotide symbols overlap with the original NTv3 alphabet, so 44 rows were added to the 11-token base vocabulary, yielding 55 model tokens; each tokenizer uses the relevant subset (Supplementary Table 1). Homozygous A, C, G and T states dominated the pretraining corpus, while heterozygous, indel and orientation marker tokens were much less frequent (Extended Data Fig. 1; Extended Data Table 4).

### 2.2 Population-scale continual pretraining yields stable diploid models

We generated reference-aligned diploid sequences from the high-coverage phased 1000 Genomes Project cohort[26, 27, 31]. Candidate intervals were selected from GENCODE[29] v47 Basic coding exons and highly conserved UCSC 100-way phastCons elements, expanded to provide language-modeling context and merged across nearby elements. The final interval set comprised 140,099 regions spanning approximately 943 Mb, or 30% of GRCh38 (Extended Data Table 1).

We initialized 8-million- and 100-million-parameter models from the corresponding Nucleotide Transformer V3[28] PRE checkpoints and resized the embedding layer for each active diploid vocabulary. Rather than initializing the new rows at random, each new heterozygous embedding was seeded from the constituent nucleotide embeddings it comprises, followed by a 5,000-step vocabulary warm-up in which the backbone was frozen and only the newly added embedding rows were updated. Diploid models and the haploid control models were then trained with masked language modeling (MLM) on 4,096-token windows; phase-aware models were additionally evaluated with Contrastive Phase Loss (CPL). A vocabulary-adapted control received the embedding warm-up but no diploid continual pretraining, and is reported as the no-continual-pretraining row of Table 1. Training and validation partitions were defined at the individual and pedigree levels to prevent close relatives from crossing splits.

**Table 1.** Compound-heterozygous *cis*-*trans* classification and phase-swap sensitivity.

| Configuration | Frozen 8M | Frozen 100M | Fine-tuned 8M | Fine-tuned 100M |
| --- | --- | --- | --- | --- |
| *Baselines* |  |  |  |  |
| No continual pretraining * | 0.512 | 0.507 | 0.506 | 0.519 |
| Haploid continuation | 0.491 ± 0.002 | 0.508 ± 0.007 | 0.533 ± 0.003 | 0.521 ± 0.008 |
| Diploid, phase removed | 0.504 ± 0.008 | 0.501 ± 0.007 | 0.519 ± 0.030 | 0.515 ± 0.010 |
| *Diploid, phase-retaining (ours)* |  |  |  |  |
| Marker-phase, MLM | 0.548 ± 0.005 | 0.540 ± 0.020 | 0.566 ± 0.023 | 0.560 ± 0.014 |
| Marker-phase, MLM + CPL | 0.555 ± 0.016 | 0.621 ± 0.005 | 0.610 ± 0.008 | 0.630 ± 0.014 |
| **Marker-phase, MLM +**<br>**CPL, full epoch primary**<br>**configuration** | **0.613±0.009** | **0.624 ± 0.005** | **0.649±0.005** | **0.639 ± 0.005** |

Both model scales trained stably, indicating that the expanded genotype-aware vocabularies could be adapted from a haploid sequence model without disrupting optimization. The haploid continual-pretraining control accounted for additional exposure to 1000 Genomes-derived sequences, allowing subsequent comparisons to separate the effect of population-scale continual pretraining from the effect of native diploid representation.

### 2.3 A compound-heterozygous benchmark isolates *cis*-*trans* phase

We next constructed the primary evaluation task around a clinically important distinction that cannot be recovered from haploid variant identity representation alone: whether two pathogenic variants in the same recessive gene occur on different homologues or on the same homologue (Fig. 1d). Under the benchmark construction, a *trans* configuration disrupts both modeled gene copies, whereas the corresponding double-*cis* configuration places both injected variants on one homologue and leaves the other without either injected variant.

The benchmark was generated by injecting curated pathogenic variants into individual 1000 Genomes haplotype backgrounds across a 117-gene panel, including 28 genes eligible for autosomal-recessive configurations. The focused *cis*-*trans* dataset contained 9,460 examples partitioned into 6,669 training, 980 validation and 1,811 test examples. Splits were family-aware and leakage-controlled: test individuals were held out, and no test variant, variant pair or *cis*-*trans* twin of a test pair appeared in training or validation.

Because this benchmark most directly tested the capability targeted by the study, its validation AU-ROC was the primary model-selection criterion. Model selection was performed on the validation partition only, with variant-effect benchmarks used as secondary safeguards against severe capability loss. The locked test partition was evaluated after configuration selection. This selection alignment is reported explicitly and is considered when interpreting the headline phase result.

### 2.4 Phase-sensitive pretraining improves configuration-aware representations

Models whose inputs do not distinguish relative phase cannot exceed chance on this task in principle, because their cis and trans examples are identical after encoding. Consistent with this, the no-pretraining control, the haploid continual-pretraining control and the unphased diploid model produced frozen-probe AUROCs of 0.491 to 0.512 and fine-tuned AUROCs of 0.506 to 0.533 across 8-million- and 100-million-parameter models (Table 1). Additional exposure to human sequence and downstream supervision did not by themselves recover the relevant haplotype arrangement.

Phase-aware tokens trained only with masked language modeling exposed orientation to the model but did not reliably force the representation to use it. Adding Contrastive Phase Loss improved frozen-probe AUROC for every phase-aware tokenizer and model-size combination, and the phase-aware Contrastive Phase Loss models outperformed all baselines by a clear margin. The strongest frozen result was obtained by the 100-million-parameter marker-phase model trained for a full epoch at 2 × 10^-4^ with Contrastive Phase Loss (0.624 ± 0.005), against 0.540 ± 0.020 for its masked-language-modelling counterpart and near-chance values for every phase-blind baseline; the same configuration reached 0.639 ± 0.005 after full fine-tuning (Table 1). Because the phase-blind conditions were evaluated on the same examples and remained near chance, this margin cannot be attributed to variant identity, allele count or gene content, and therefore reflects recovered haplotype configuration. These are point estimates from a synthetic, task-aligned benchmark and do not establish superiority to a two-stream haplotype encoder.

Representation-level analysis supported the same interpretation. Masked language modeling alone produced high cosine similarity between original heterozygous contexts and the same contexts after a global exchange of the two homologue labels, indicating that orientation was present in the input but largely ignored by the representation. Contrastive Phase Loss sharply reduced this similarity and increased sensitivity to phase reversal (Supplementary Table 6). This is expected, since the cosine quantity is the Contrastive Phase Loss objective itself evaluated on held-out data, and it is therefore a manipulation check rather than independent evidence. A global exchange of homologue labels also does not change whether two variants are in cis or in trans, so the analysis is a diagnostic of orientation sensitivity, not a biological phase endpoint; the paired cis-trans benchmark provides the evidence for relative-phase discrimination. The no-projector condition produced stronger separation still under this diagnostic, but the projector-based loss was selected as the more conservative general-purpose configuration.

Increasing parameter count did not yield consistent gains across tokenizers or endpoints. In several comparisons the 8-million-parameter model exceeded the corresponding 100-million-parameter model, whereas the reverse held elsewhere. Model scale is therefore treated as an empirical factor rather than an assumed source of monotonic improvement.

The two phase-retaining tokenizers showed different trade-offs. The ordered-phase tokenizer provides a compact single-token-per-position representation, whereas the marker-phase tokenizer, which separates unordered allele content from an explicit orientation marker, achieved the strongest compound-heterozygous frozen probe and more consistently preserved indel performance (Supplementary Table 7). We therefore carried the marker-phase tokenizer with projector-based Contrastive Phase Loss forward as the primary configuration, and retained the ordered-phase model as a useful alternative where a compact representation is preferred. These observations support the choice of configuration for this study but do not establish one tokenizer as universally preferable.

### 2.5 Generalization across zygosity, short indels and population structure

We next evaluated zygosity-sensitive ClinVar SNVs, zero-shot ClinVar indels and general variant-effect controls (Fig. 3; Table 2). These tasks probe different properties and are interpreted separately; they are not combined into a claim that every diploid model condition improves every genomic prediction task.

**Table 2.** ClinVar variant-effect prediction by input encoding.

| Configuration | Diploid | Haploid | Biallelic | $\Delta$ diploid – haploid |
| --- | --- | --- | --- | --- |
| **8 million parameters (NTv3_8M_pre)** |  |  |  |  |
| Haploid continuation | NA | 0.812 | 0.783 | NA |
| Diploid, phase removed | 0.853 | 0.698 | 0.723 | +0.155 |
| Marker-phase, MLM | 0.836 | 0.842 | 0.738 | −0.006 |
| Marker-phase, MLM + CPL | 0.798 | 0.660 | 0.573 | +0.138 |
| **Marker-phase, MLM + CPL, full epoch primary configuration** | **0.852** | **0.819** | **0.818** | **+0.033** |
| **100 million parameters (NTv3_100M_pre)** |  |  |  |  |
| Haploid continuation | 0.791 | 0.661 | 0.583 | +0.120 |
| Diploid, phase removed | 0.830 | 0.660 | 0.622 | +0.170 |
| Marker-phase, MLM | 0.757 | 0.696 | 0.715 | +0.061 |
| Marker-phase, MLM + CPL | 0.746 | 0.592 | 0.616 | +0.154 |
| **Marker-phase, MLM + CPL, full epoch primary configuration** | **0.755** | **0.541** | **0.555** | **+0.214** |

ClinVar supervised variant-effect prediction; test AUROC, mean over three probe seeds. Chance = 0.5. Each trained model is read out three ways on an identical variant set (the diploid tokenization, a single haplotype, and both haplotypes as two separate sequences), so the comparison isolates the input encoding rather than the pretraining recipe. Across the eleven arms evaluated the diploid readout is highest in 10 of 11; mean paired advantage over the haploid readout 0.106 AUROC, Wilcoxon signed-rank exact $P = 0.0029$ , $n = 11$ arms with probe seeds averaged before pairing. On Matthews correlation the diploid readout is ahead in 11 of 11 (mean advantage 0.117, $P = 0.00098$ ). The single exception is the 8M marker-phase MLM arm, where the haploid readout leads by 0.006.

The mode-of-inheritance-aware ClinVar SNV benchmark asks whether a variant, in a given zygosity, is pathogenic. This is a per-site question about allelic dosage: it does not require knowing which haplotype carries the variant. Under the two-channel account developed above we therefore expect the diploid encoding to help and the phase objectives not to, and this is what we observe (Fig. 3a). The highest score was obtained by the phase-removed control (0.853 at 8 million parameters) (Table 2), a condition that retains diploid dosage while discarding phase entirely, and it exceeded the haploid continuation (0.812), which discards both. Marker-phase CPL models performed comparably but not better, reaching 0.798 at 8 million parameters and 0.852 after full-epoch training. Results were strongly scale-dependent in a direction we cannot presently explain: the corresponding 100-million-parameter models reached only 0.746 and 0.755 and did not exceed the strongest controls.

Because the aggregate task mixes pathogenicity, inheritance category and zygosity, it does not by itself isolate dosage. The within-model encoding comparison in Table 2 is the analysis that does, and it shows the diploid readout ahead of the haploid readout in 10 of 11 arms by a mean of 0.106 AUROC including in the conditions trained without any phase objective.

On the held-out ClinVar indel benchmark the highest AUROCs occurred in phase-blind conditions: 0.727 for marker-phase MLM at 8 million parameters and 0.728 for the phase-removed control at 100 million, with every CPL condition lower at the same scale (Table 3; Fig. 3).

**Table 3.** ClinVar indel pathogenicity prediction by input encoding.

| Configuration | Diploid | Haploid | Biallelic | $\Delta$ diploid – haploid |
| --- | --- | --- | --- | --- |
| **8 million parameters (NTv3_8M_pre)** |  |  |  |  |
| Haploid continuation | NA | 0.550 | 0.218 | NA |
| Diploid, phase removed | 0.705 | 0.540 | 0.214 | +0.165 |
| Marker-phase, MLM | 0.727 | 0.546 | 0.217 | +0.181 |
| Marker-phase, MLM + CPL | 0.641 | 0.511 | 0.200 | +0.130 |
| **Marker-phase, MLM + CPL, full epoch primary configuration** | **0.658** | **0.516** | **0.202** | **+0.142** |
| **100 million parameters (NTv3_100M_pre)** |  |  |  |  |
| Haploid continuation | NA | 0.529 | 0.205 | NA |
| Diploid, phase removed | 0.728 | 0.545 | 0.213 | +0.183 |
| Marker-phase, MLM | 0.708 | 0.536 | 0.208 | +0.172 |
| Marker-phase, MLM + CPL | 0.706 | 0.509 | 0.195 | +0.196 |
| **Marker-phase, MLM + CPL, full epoch primary configuration** | **0.717** | **0.539** | **0.214** | **+0.178** |

ClinVar zero-shot indel pathogenicity prediction, sequence embedding L2 AUROC. The mean paired advantage is 0.154 AUROC (range 0.084 to 0.196); Mean AUROC is 0.679 for diploid, 0.524 for haploid, and 0.206 for biallelic input.

In the indel task, biallelic mode, where a genotype’s two alleles are compared to each other rather than to the reference, we hypothesize that the low embedding-distance AUROC (0.20 cosine; 0.21 for L2), varying by less than 0.03 across fifteen checkpoints spanning 8M–100M parameters and three pretraining objectives, is a structural consequence of the comparison: for a homozygous genotype the two alleles are identical strings, so their distance is exactly zero irrespective of sequence content, and the score reduces to a heterozygosity indicator. Because the benchmark labels heterozygous recessive genotypes benign and their homozygous counterparts pathogenic, and benign variants comprise a small minority of the set, such an indicator is anti-correlated with the label; a simulated zygosity-only score reproduced both the observed AUROC and its below-prevalence AUPRC. Padding both alleles to equal length, which removes indel size as a covariate, left the metric unchanged or marginally lower, confirming that it does not read sequence content. Biallelic comparison therefore quantifies zygosity rather than variant effect, and the haploid comparison is the appropriate contrast for this benchmark.

### 2.6 Preservation of conventional haploid variant-effect capability

BRCA1 and TraitGym variant effect prediction tasks, in addition to general haploid evaluation tasks in the GFMBench-API suite[30] were used as preservation controls as their supplied sequences are haploid and their labels do not require zygosity or phase. After mapping each nucleotide to its corresponding homozygous diploid state, performance was better in most tasks and on-par or slightly lower in the rest across model conditions (Extended Data Table 3; Supplementary Table 4). These controls show that vocabulary adaptation did not systematically erase ordinary haploid genomic representation information.

### 2.7 General haploid capabilities are broadly preserved after diploid continual pretraining

To test whether the expanded vocabulary and the phase objective degrade capabilities that have nothing to do with diploidy, we evaluated frozen representations on three tasks from the Genome Understanding Evaluation suite (promoter identification, splice-site reconstruction and transcription-factor binding) using a linear probe over sequence embeddings at 300 bp (Supplementary Table 8). These tasks contain no phase information: GUE sequences are haploid reference DNA and encode to the five base nucleotide rows under every tokenizer in this study, so the 44 diploid-specific rows are never instantiated and all conditions receive identical input tokens. Any difference is therefore attributable to the backbone rather than to the encoding. At 8 million parameters, every continually pretrained condition improved over the original model on every task,raising mean Matthews correlation from 0.454 to between 0.601 and 0.610. At 100 million parameters, gains were smaller and task-dependent. The haploid continuation, the phase-removed diploid model and the marker-phase model trained with masked language modeling each fell at most 0.010 MCC below the original checkpoint on transcription-factor binding, and the phase-removed model fell 0.008 below on promoter identification. Marker-phase models trained with CPL were at or above the original checkpoint on all three tasks (0.815, 0.560 and 0.587; mean 0.654 against 0.609). Diploid continual pretraining therefore does not cost general sequence modelling on these tasks. Because GUE comprises three tasks under a single probe, it is reported as a capability control rather than as evidence of improved general capability; the broader comparison against the unmodified backbone is given in Supplementary Table 4.

### 2.8 CFTR examples illustrate genotype-dependent model behavior

As a clinically familiar case study, we examined zero-shot behavior in Cystic Fibrosis Transmembrane Conductance Regulator (CFTR), (Fig. 4). Across ClinVar-annotated CFTR SNVs, reference-versus-alternative log-likelihood-ratio scores separated pathogenic from benign variants with AUROC 0.780. The nonsense variant G542X fell within the pathogenic score range, whereas the variable-penetrance missense variant R117H received a lower, benign-like score.

**Figure 4.**
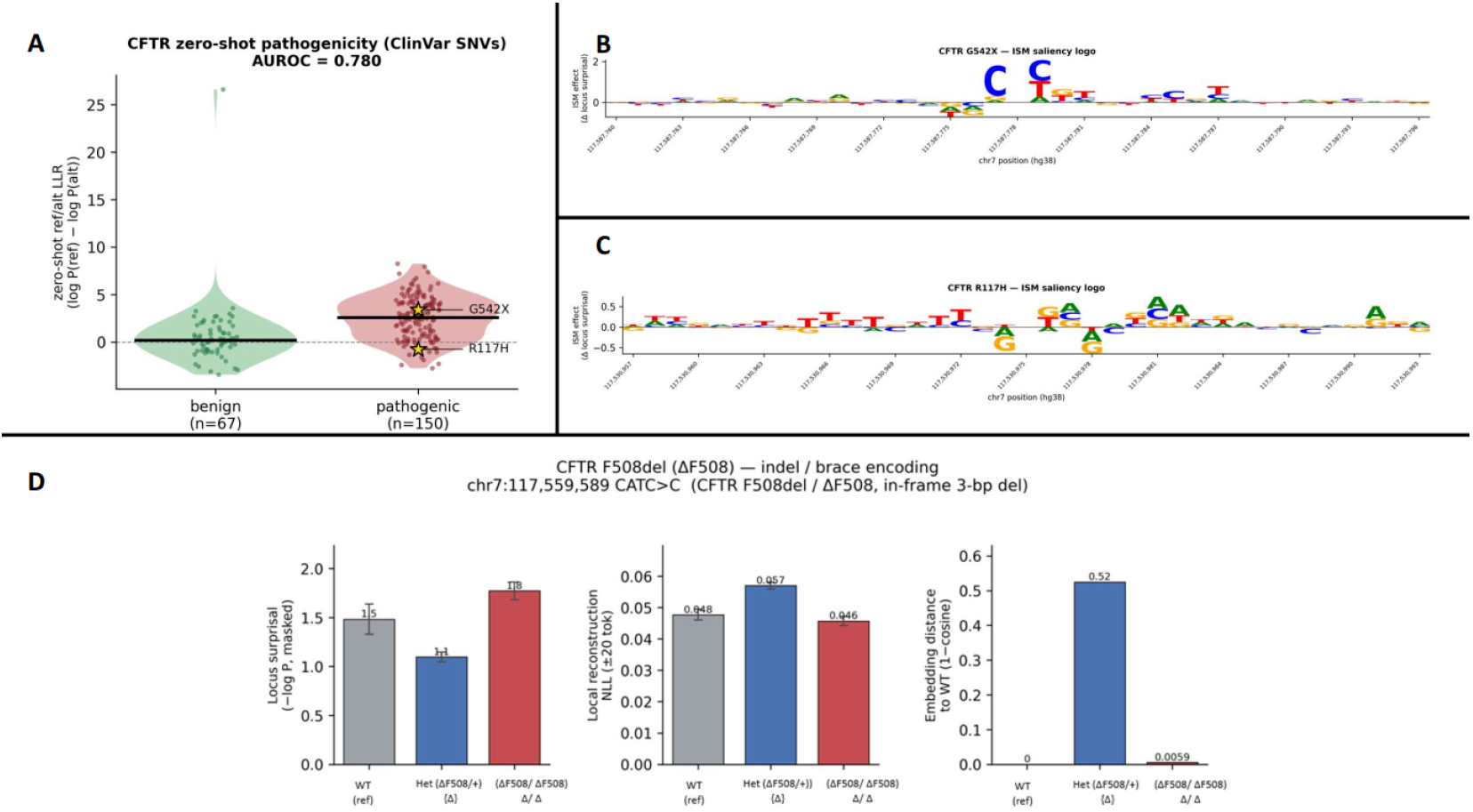
CFTR case study and broader capability controls. a, Violin plot of zero-shot reference-versus-alternative log-likelihood-ratio scores for ClinVar-annotated CFTR SNVs (benign n = 67, pathogenic n = 150; AUROC 0.780), with G542X and R117H marked. b, In-silico mutagenesis sequence logo for G542X, showing a high, localized variant-centered signal. c, Sequence logo for R117H, showing a weaker and more diffuse local signal. d, Genotype-level local scores for F508del in wild-type, heterozygous and homozygous states, showing a distinct heterozygous-deletion signal relative to wild type.

In-silico mutagenesis produced a localized attribution pattern around G542X and a weaker, more diffuse pattern around R117H, indicating that the local score is driven by variant-local sequence evidence rather than gene-level background alone. For the canonical F508del deletion, heterozygous and homozygous genotypes produced distinct local score profiles, providing an example in which the diploid representation responds differently to two dosage states of the same indel.

While these examples are descriptive and do not establish clinical validity, they show that the model can yield variant-local and genotype-dependent signals in a well-characterized recessive disease gene. They do not test diagnostic accuracy, penetrance or patient-level outcomes.

## 3 Discussion

This study identifies genotype configuration as a missing representational variable in most sequence-based genomic foundation models. We address this gap by encoding both homologues within one reference-aligned stream, so the proposed tokenizers make zygosity, allele dosage, explicit indel sequence and optional phase orientation available before downstream aggregation. The continually pretrained models demonstrate that this expanded vocabulary can be learned from population-scale human genomes, while the haploid continual-pretraining control separates native diploid representation from the effect of additional human-sequence pretraining.

The controlled compound-heterozygous benchmark is central to this conclusion. Generic pathogenicity tasks can often be solved from variant identity, conservation or molecular consequence without representing the arrangement of alleles. Here, *cis* and *trans* examples contain the same pathogenic pair and differ only in haplotype configuration. The near-chance performance of models without phase information, and the clear margin achieved once phase-sensitive pretraining is applied, indicate that the benchmark is probing the intended representational property. The benchmark implementation and fixed split logic provide a reusable framework for evaluating future phase-aware models.

The work complements, rather than replaces, previous strategies for modeling human variation. Genotype-imputation models learn haplotype and linkage-disequilibrium structure over variant panels[19, 20], and dual-allele sequence approaches can preserve zygosity by encoding the two homologues separately[24, 25]. Our approach instead creates one sequence-native representation at each aligned locus and supports explicit indel alleles. This design removes the need for two encoder passes and gives the model direct access to allele pairing, but it also imposes a reference-aligned representation and expands the vocabulary. Which representation is preferable will depend on the downstream task, context length and available phasing information.

The clinical relevance lies in the class of questions the representation can express. Carrier status, biallelic disease, hemizygosity and compound heterozygosity cannot be reduced to an isolated alternative allele. A genotype-aware model could eventually support inheritance-constrained prioritization, ranking of candidate variant pairs and integration of phase into rare-disease workflows. The present study does not demonstrate those uses. The ClinVar labels apply simplified inheritance rules, the cis-*trans* benchmark is computationally generated and the CFTR analyses are illustrative. Accordingly, the appropriate claim is that the framework represents configurations relevant to clinical interpretation, not that it performs clinical diagnosis or replaces expert variant assessment.

Contrastive Phase Loss also reveals an important modeling distinction. Supplying phase-aware tokens is not equivalent to learning a phase-sensitive representation: masked language modeling can largely ignore orientation when it is weakly predictive of the pretraining objective. The contrastive objective increased sensitivity to phase reversal and improved *cis*-*trans* discrimination, but the strongest separation did not always preserve other capabilities, which motivated selection of the projector-based formulation. Together with the broadly comparable general variant-effect controls, this argues against presenting diploid pretraining as a universal performance improvement. It is better understood as a targeted intervention that supplies the information required by configuration-dependent tasks without systematic degradation on the controls tested.

Several limitations define the next steps. First, the compound-heterozygous examples are synthetic and use curated variants injected into population haplotypes; naturally observed affected individuals may differ in transcript choice, variant severity, penetrance, modifiers, ancestry and ascertainment. Second, statistically phased population data contain errors, particularly in complex or poorly tagged regions. Trio-based, long-read and haplotype-resolved data should provide stronger training and evaluation substrates. Third, although the NTv3 backbone supports much longer native contexts, this study used 4,096-token windows and therefore primarily captures local configuration rather than many gene-scale or regulatory relationships. Fourth, the tokenizer currently focuses on SNVs and short indels rather than structural variants, copy-number changes, repeat expansions, inversions or complex rearrangements. Fifth, ClinVar and HGMD are enriched for well-studied genes, severe Mendelian phenotypes and historically overrepresented populations. Sixth, all model experiments use the NTv3 backbone, so architectural generalization to other transformers, state-space models or long-context backbones remains untested. Finally, the primary phase task guided model selection and ancillary benchmarks were used as guardrails; independent locked cohorts and external task families are therefore needed to establish generalization.

Together, the tokenization schemes, pretrained models and benchmark suite provide a practical foundation for genotype-configuration-aware genomic modeling. The results show that phase-dependent distinctions absent from haploid or unordered inputs can become recoverable when both the representation and the pretraining objective are designed for them. The next test is not another synthetic benchmark alone, but evaluation in naturally phased clinical cohorts with molecularly confirmed diagnoses and realistic inheritance-aware endpoints.

## 4 Methods

### 4.1 Reference-aligned diploid encoding and terminology

The diploid tokenizer maps the allelic state of the two homologues at each reference-aligned position to one token. The term encoding refers to the overall representation; tokenizer refers to the deterministic mapping procedure; vocabulary refers to the available symbols; and model condition refers to a tokenizer, backbone and training objective. Symmetric states AA, CC, GG and TT are encoded as A, C, G and T. The unphased tokenizer uses one symbol for both reciprocal heterozygous orders. The ordered-phase tokenizer uses separate symbols for reciprocal orders. The marker-phase tokenizer combines the unphased allele-pair symbol with <DIR_FWD> or <DIR_REV>.

All active tokenizers include <SNP>, <INDEL_SPAN_START> and <INDEL_SPAN_END> metadata tokens. Single-allele symbols are defined in the union vocabulary for future hemizygous encoding, but they were not active in the reported corpus. In the current implementation, male X-linked examples were rendered as homozygous states and therefore do not provide evidence that the model explicitly represents hemizygosity. The exact ordered-pair and marker mappings are fixed by the released token table (Supplementary Table 1).

### 4.2 Reference-aligned SNV and indel encoding

For an SNV, the allele pair is encoded at the reference coordinate and preceded by <SNP>. For a short indel, each homologue is rendered locally with explicit span boundaries; the two allele strings are aligned relative to a shared left anchor; and the shorter allele is padded with gap states. Each aligned nucleotide or nucleotide-gap pair is then converted to one diploid token (Fig. 2b).

Reference bases retain GRCh38 coordinates. Inserted bases receive ordered offsets between adjacent reference coordinates, allowing the token stream to be mapped back to the reference. The coordinate metadata was not supplied to the model. The implementation accepts explicit A/C/G/T SNVs and short indels. Symbolic alleles, structural variants, copy-number variants and complex rearrangements were excluded.

### 4.3 Construction of the pretraining corpus

Phased variants were obtained from the high-coverage 1000 Genomes Project release comprising approximately 3,200 individuals, including 602 complete trios[31]. We partitioned the cohort into *training* (2,560 samples, 79.95%) and *test* (642 samples, 20.05%) subsets under two competing constraints: preventing information leakage between relatives and preserving demographic representativeness. The first is binding, and where the two conflict it takes precedence. Because family members share large fractions of their genomes through Mendelian inheritance, any relative appearing in both subsets would allow a model to exploit familial variant patterns rather than generalizable genomic features, inflating apparent performance; we therefore treated the connected pedigree, not the individual, as the indivisible unit of assignment. Delimiting those pedigrees was not trivial: six biologically connected families span multiple FamilyIDs, and partitioning on FamilyID directly separated related individuals in two cases. We instead reconstructed pedigrees with a union-find procedure that links each individual to its parents and to full and half-siblings, yielding 2,001 connected components (122 more than the 1,879 FamilyID groups), ranging from single unrelated individuals (1,407 components) to six-member three-generation pedigrees. This variability in unit size introduces a second difficulty, since allocating by component count rather than sample count would distort the sample-level ratio, and larger components carry disproportionately many parents and children, skewing trio-role composition. We therefore stratified assignment by superpopulation × component size, allocating whole components so as to approximate 80% of samples within each stratum while retaining at least one component in each subset wherever a stratum contained two or more. Superpopulation was stratified explicitly because allele frequencies and linkage-disequilibrium structure are population-specific, whereas sex is balanced indirectly through component assignment, trio role through the component-size stratification, and 26-population membership through the superpopulation stratification; none of these three is explicitly constrained. Assignment used a fixed random seed (42) for all shuffling. The resulting partition leaves zero pedigrees straddling the train/test boundary and deviates ≤0.6% from the 80/20 target across superpopulations and <=0.2% across sex and trio role; residual per-population deviation (mean 3.7%, maximum 10.9% for PUR) reflects the discreteness of indivisible family units within small strata, as does the imbalance in the rarest component-size strata, where the single two-member and four six-member components could not be divided near 80/20.

Chromosome-level phased VCF files were converted to indexed BCF files to enable efficient sample-wise processing, extracted per individual, and merged with UCSC hg38 to produce two haplotypes and the corresponding reference-aligned diploid-token sequence (Extended Data Fig. 2).

To focus pretraining on functional and evolutionarily constrained sequence, candidate regions were drawn from GENCODE v47 Basic coding exons and UCSC 100-way vertebrate phastCons conserved elements with log-odds score >500, yielding approximately 336,500 elements spanning approximately 217 Mb (about 7% of hg38). A filter-inflate-merge pass removed elements shorter than 10 bp, expanded each retained element to a minimum 4,094-bp context window and merged windows separated by 300 bp or less. Only canonical chromosomes 1-22, X and Y were retained. The resulting 140,099 intervals covered approximately 943 Mb, or about 30% of GRCh38 (Extended Data Table 1). In sequence without indels, one reference-aligned base generally yields one diploid state token; SNV and indel markers, gap states and inserted bases add tokens, so base-pair and token counts are not always identical. All sequences were then processed with the tokenization scheme described above.

#### Window construction

Diploid sequences were rendered for each of 2,560 training individuals across the 140,099 intervals described above. Sequences identical across individuals were deduplicated, leaving 110,887,674 records. Each record was windowed at 4,096 tokens with stride 3,686, giving 228,645,057 windows (2.1 per record), 94.7% of which contained at least one variant. These were subsampled to all variant-bearing windows, all GRCh38 reference windows and a random 40% of the remainder, giving 221,431,899 training windows. Reference windows (0.09% of the index) were sampled at twice their base rate. One epoch was 221.3 million samples drawn with replacement, or 922,206 optimizer steps at an effective batch size of 240. Record frequency in the raw data, recorded during deduplication, was applied as a per-sample loss weight capped at 20.

#### Marker dropout

To help with robustness in clinical settings with noisy phased reads, the SNV anchor token was randomly deleted with probability 0.15 during training.

#### Model initialization and vocabulary adaptation

Models were initialized from the InstaDeepAI NTv3_8M_pre and NTv3_100M_pre checkpoints. NTv3 takes single-nucleotide input, so each diploid state can be registered as one additional codepoint. We reserved 44 rows spanning the union of diploid symbols across the three tokenizers, giving a 55-token vocabulary, and resized the input embedding and language-model head to match. Any single corpus instantiates a subset of these: the marker-phase corpus contains 20 distinct symbols, 15 of them diploid-specific: ten allele-pair and gap symbols, the SNV marker, the two indel-span delimiters and the two orientation markers. Reserved rows for symbols absent from a given corpus are never instantiated and receive no input-side gradient.

New tokens were seeded from the nucleotides they compose: a heterozygous token took the mean of its two parent embeddings, phase-ordered symbols were offset by ±0.05 of the parent difference so reciprocal pairs stay separable, and single-parent symbols were cloned from their parent. All seeded tokens received Gaussian noise (s.d. 0.03). The head’s 44 new tokens are tied to the mean of the pretrained head rows.

The new tokens were warmed up for 5,000 steps with the backbone frozen with the pre-existing nucleotide token embeddings held static during the warmup. Warm-up used 1,024-token sized variant-containing windows, peak learning rate 1 × 10^-4^ and weight decay 0.01 (Supplementary Table 2) . One warm-up was run per tokenization scheme and model size, and all downstream runs resumed from it.

### 4.4 Continual pretraining

Diploid and control haploid models were continually pretrained with a masked language modeling objective rather than trained from random initialization. At each step, each eligible token position was independently selected for masking with probability 0.15. Special tokens and the homozygous-N symbol were excluded from selection before sampling. Of the selected positions, 80% were replaced with the [MASK] token, 10% were replaced with a token drawn from a frequency weighted distribution. The 44 non-special, non-marker vocabulary rows, and 10% were left unchanged. This is the standard BERT-style corruption policy. The 10%-random and unchanged components prevent the encoder from assuming that an observed token is necessarily unmasked and correct, which matters more here than in haploid modeling because an unmasked heterozygous (or SNP) symbol is itself a strong, locally informative signal, so the encoder must not treat an observed token as necessarily correct.

Masking was applied to whole diploid tokens rather than independently to the two nucleotides. Masking a heterozygous position therefore hides the joint allelic state of both homologues. Recovery requires predicting zygosity and, for phase-aware tokenizers, orientation, rather than prediction of one nucleotide alone.

Production runs used 4,096-token windows, global effective batch size of 240, bfloat16 precision, fused AdamW (beta1 = 0.9, beta2 = 0.98), peak learning rate 2 x 10^-4, 10,000 warm-up steps with linear decay, weight decay 0.005 excluding normalization, bias and embedding parameters, and a maximum gradient norm of 1.0. A matched run at 5 x 10^-5 is reported as a learning-rate ablation (Supplementary Table 3) . Training was performed on an NVIDIA DGX system containing eight H100 SXM Tensor Core GPUs. Marker dropout was 0.15. The full run comprised 922,206 optimizer steps (one epoch) over the 110.9 million training records of the phased cohort; Supplementary Table 2 provides the complete configuration.

Haploid controls used the same initialization and continual-pretraining framework but encoded each genome as a single haploid sequence rather than a diploid token stream, and were trained with masked language modeling only. Haploid controls were trained at the 60,000-step ablation budget.

### 4.5 Contrastive Phase Loss (CPL) as an orientation-sensitive auxiliary objective

Phase-aware diploid tokens expose haplotype orientation to the model, but masked language modeling does not explicitly reward representations that change when the encoded orientation is reversed.

Empirically, opposite orientations of the same heterozygous allele pair, such as C/A and A/C, produced highly similar representations, indicating that orientation was available in the input but largely ignored by the encoder. This matters most for compound-heterozygous configurations, where two pathogenic variants in *trans* can disrupt both modeled gene copies, whereas the same variants in *cis* may leave one haplotype intact.

We therefore introduced a Contrastive Phase Loss (CPL). CPL was applied only to diploid models whose inputs retained phase orientation, that is, the ordered-symbol tokenizer and the unphased-symbol-plus-directional-marker tokenizer. For each phase-aware training window we construct an orientation-reversed counterpart by inverting the encoded haplotype orientation at every heterozygous position: either by flipping the directional marker (</>) or by replacing each ordered symbol with its reverse-orientation counterpart (for example Á <-> Ç). All non-phase tokens and the masking pattern are held fixed, so the only difference between the two views is the encoded orientation.

Let h_i_ and ̃h_i_ denote the hidden states at heterozygous position i in the original and phase-reversed sequences, and let g(.) denote the projection head. H is the set of heterozygous positions shared by the paired views, and |H| is its cardinality. The auxiliary contrastive term minimizes cosine similarity between projected hidden states at corresponding positions. Phase reversal is therefore encouraged to produce distinct and, at the isolated optimum, anti-aligned representations in the projection space, making the supplied orientation signal accessible to downstream models. This term is optimized jointly with masked language modeling, which maintains pressure to preserve sequence-predictive information while the projection head captures phase sensitivity. The final objective is shown below:

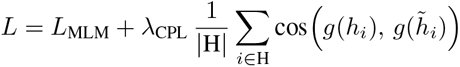

λ was selected by a sweep over {0, 0.25, 0.5, 1.0} together with two two-stage schedules in which CPL was switched on or off at a checkpoint (Supplementary Table 5). Compound-heterozygous discrimination was insensitive across the constant-weight range (0.596 to 0.616) and phase sensitivity saturated throughout, so λ = 0.5 was taken as the midpoint of a flat region. Both two-stage schedules markedly weakened phase sensitivity without improving discrimination, so a constant weight was retained. Projector and no-projector variants were evaluated. The selected formulation used a 64dimensional projector (Table S6). Without a projection head the objective is minimised by a global sign inversion of the representation under phase reversal, and the no-projector variant does exactly this, reaching a cosine similarity of −0.999. That solution satisfies the loss without encoding phase where it is useful: at 100 million parameters it scored 0.532 against 0.586 on frozen compound-heterozygous discrimination and 0.767 against 0.815 on the seven-scenario zygosity task. Interposing a learned projection prevents the objective from being discharged by such a reflection, so phase sensitivity must be carried by the backbone representation itself.

### 4.6 Ablation design and model selection

Ablations compared no-pretraining controls, haploid continual-pretraining control, unphased diploid models, phased ordered-symbol models and unphased-plus-directionality models, with and without Contrastive Phase Loss. Both model sizes were trained for 60,000 ablation steps from the shared vocabulary-warmed checkpoint, corresponding to 14.4 million training windows at effective batch size 240. Runs used 4,096-token windows, 1,000 warm-up steps.

The ablation addressed three questions. First, haploid and diploid continual pretraining separated the effect of native genotype representation from additional exposure to population-scale human sequence. Second, comparison of unphased and phase-retaining tokenizers tested whether preserving haplotype order improved downstream performance. Third, comparison with and without CPL tested whether explicit phase-sensitive supervision improved *cis*-*trans* discrimination. A biallelic baseline was included at benchmarking time using haploid-model representations rather than a separately pretrained biallelic model; it approximates diploidy downstream but never processes a unified diploid genotype during pretraining. Together, these comparisons isolate additional human-genome pretraining, native representation of both alleles and phase-sensitive learning as candidate sources of gain.

The primary selection criterion was validation AUROC from a frozen linear probe on the compound-heterozygous cis-trans task, averaged over three probe seeds. Variant-effect tasks were used as secondary guardrails against severe degradation. The selected configuration was then continued to full pretraining. The haploid control was not continued to full pretraining; exposure-controlled comparisons are therefore reported at the 60,000-step budget. The compound-heterozygous test split was not used for selection, but the task family was selection-aligned; this is reported as a limitation.

### 4.7 Evaluation modes and benchmark implementation

Benchmark execution was standardized through GFMBench-API^30^. The ClinVar diploid SNV task pairs a variant and reference sequence per record with a binary pathogenic/benign label, split by chromosome with chromosomes 19–22 held out for test and no validation split; the ClinVar indel task follows the same interface with sequences rendered per haplotype. The three input modes do not evaluate identical row sets. Diploid and biallelic modes emit one record per variant, whereas haploid mode expands each heterozygous variant into two records, an ALT haplotype and a REF haplotype, both scored against the same reference, so the haploid row set is larger and its label prevalence differs. Differences between haploid and diploid scores therefore reflect both the model input and the evaluation sample, and within-mode comparisons are the interpretable ones.

Three input modes were compared for variant-centric tasks (Fig. 1c). Diploid mode encodes the realized genotype in one sequence-native stream and scores it in a single forward path. Haploid mode encodes the two alleles separately, scores REF and ALT for each allele, and averages the resulting predictions downstream. Biallelic mode supplies allele 1 and allele 2 simultaneously without REF/ALT substitution semantics, but uses haploid-model representations rather than a separately pretrained biallelic model; it is our reimplementation of the dual-allele strategy of refs Saadat et al. and Rashidy et al.[24, 25], which is the closest available comparison because the SNP2Vec-family checkpoints are not publicly released. Because these modes differ in their available input and, for some zero-shot analyses, in their optimal scoring rule, comparisons are interpreted accordingly.

### 4.8 Compound-heterozygous benchmark

Curated pathogenic variants from HGMD Professional 2023.2[32] were injected into 1000 Genomes haplotype backgrounds across a 117-gene panel, 28 genes of which were eligible for autosomal-recessive configurations.

Each sequence window was 6,144 bp. Two datasets were built from the same generator. The first spans seven inheritance and zygosity scenarios: autosomal-dominant heterozygous, autosomal-recessive carrier, autosomal-recessive homozygous, autosomal-recessive compound heterozygous in *trans*, double heterozygous in *cis*, X-linked-recessive female carrier and X-linked-recessive male hemizygous, and is close to saturated after fine-tuning. The second is restricted to the compound-heterozygous contrast alone and is substantially harder: measured on the same models, fine-tuning reaches 0.813 to 0.989 AUROC on the seven-scenario task against 0.506 to 0.649 on the compound-heterozygous contrast (Extended Data Table 2). This is the only contrast that cannot be resolved from variant identity, gene identity or allele count, and therefore the only one requiring the model to represent which haplotype each variant lies on.

That contrast is the primary task: compound-heterozygous *trans* versus double-*cis*, built by injecting two pathogenic variants that differ only in whether they lie on different haplotypes (disease-causing under a recessive model, because both gene copies are disrupted) or on the same haplotype (leaving one modeled gene copy without either injected variant). The final dataset contained 9,460 examples divided into 6,669 training, 980 validation and 1,811 test examples. The split excluded overlap of individuals, injected variants, variant pairs and *cis*-*trans* twins between the test split and either the training or the validation split, so the test phase labels cannot be seen during training or model selection.

Sequences were encoded with the diploid directional tokenization. For each example, hidden states at the two injected variant positions were extracted as e_A and e_B. The classifier input concatenated e_A, e_B, e_A − e_B and the element-wise product e_A ⊙ e_B; the difference and product terms render the same-versus different-haplotype comparison linearly separable. Frozen evaluation trained only a linear classifier on the pooled feature, measuring the phase information already present in the representation; full fine-tuning updated the backbone and classifier jointly, measuring the phase discrimination attainable after adaptation. Models were selected by validation AUROC and reported by test AUROC averaged over three random seeds. Robustness analyses compared pooling at the two known variant positions with mean-pooling across all variant positions, and evaluated a phase-swap augmentation that flips each example’s phase markers together with its label to remove any residual variant-identity cue.

The pretraining train/validation split was pedigree-aware as described above. The compound-heterozygous benchmark used the separate three-way, leakage-controlled train/validation/test partition reported in the preceding paragraph; the earlier approximately 80:20 split does not apply to this benchmark.

### 4.9 Mode-of-inheritance-aware ClinVar SNV benchmark

The SNV benchmark used the January 2026 ClinVar[33] variant summary and VCV XML releases restricted to GRCh38 germline SNVs on canonical chromosomes with a review status of at least two gold stars (multiple concordant submitters, expert-panel or practice-guideline assertions). Benign and likely benign variants were assigned variant label 0; pathogenic and likely pathogenic variants were assigned variant label 1; conflicting and uncertain records were removed. Mode of inheritance was normalized to a dominant or recessive class from the VCV records (autosomal- and X-linked-dominant mapped to dominant; autosomal- and X-linked-recessive mapped to recessive), and variants without a resolvable class were removed.

Each retained variant was instantiated in heterozygous and homozygous states. Benign variants received a negative genotype-level label in both states; pathogenic dominant variants received a positive label in both states; pathogenic recessive variants received a negative label in the heterozygous state and a positive label in the homozygous state. These synthetic outcomes are benchmark labels rather than revisions to ClinVar classification; a pathogenic recessive variant remains pathogenic when present in a heterozygous carrier. The construction is intentionally zygosity-sensitive: heterozygous and homozygous instances derived from the same recessive variant receive different labels, so variant identity alone is insufficient to determine the assigned outcome. The labels omit penetrance, variable expressivity, hypomorphic alleles, compound heterozygosity and gene-specific exceptions.

Examples comprised 1,024-bp windows centered on the variant, presented as a variant/reference sequence pair (or as allele-1/allele-2 inputs in the biallelic configuration). Chromosomes 19-22 formed the test set; the remaining autosomes and X formed the training set, so no genomic position is shared across the split. The reported distribution was 140,972 negative and 6,922 positive training instances and 21,414 negative and 976 positive test instances. Because positive prevalence is approximately 5%, training used inverse-frequency class rebalancing (a weighted random sampler with power = 1), and both AUROC and AUPRC are reported. The benchmark is evaluated in a supervised setting (fine-tuning or frozen linear probing) across the diploid, haploid and biallelic input representations.

### 4.10 ClinVar indel benchmark

The indel benchmark applied the same ClinVar release, assembly, germline filter, two-star review-status threshold and mode-of-inheritance relabelling as the SNV benchmark, but retained ClinVar insertion, deletion and indel types. Alleles were quality-filtered to explicit A/C/G/T strings (symbolic, missing and star alleles were rejected); reference and alternate alleles were required to differ in length and to share a common left-anchor base, consistent with left-aligned VCF representation, with the length change determining the insertion or deletion class. Each genotype was rendered through the local biallelic alignment and brace-delimited diploid indel encoding described above.

The benchmark contained 939 benign and 2,065 pathogenic variants, evaluated as a single held-out set with no fine-tuning. Variant effect was scored both by the cosine distance between variant and reference embeddings and by a masked log-likelihood ratio, the latter being the more appropriate score for the directional-marker tokenizer.

### 4.11 General haploid controls and CFTR analysis

All benchmark tasks, including BRCA1 and TraitGym, were executed through GFMBench-API[30]. BRCA1 and TraitGym were retained as zero-shot general-capability controls because their labels and available input sequences are haploid and do not require genotype configuration. For these tasks, each nucleotide was mapped to the corresponding homozygous diploid state (for example, A to AA), allowing the diploid encoder to process the sequence without introducing heterozygosity or phase.

For the CFTR case study, ClinVar-annotated SNVs were scored by reference-versus-alternative log-likelihood ratio. In-silico mutagenesis was used to visualize local attribution around G542X and R117H. F508del was rendered in wild-type, heterozygous and homozygous genotype states and compared using local likelihood and embedding-distance profiles. The analysis was descriptive; no patient-level clinical outcome was predicted.

### 4.12 Data availability

The pretraining corpus was derived from the publicly available high-coverage phased 1000 Genomes Project release[26, 27, 31]. Reference sequence was taken from UCSC^1^ and annotations were obtained from GRCh38, GENCODE v47 and UCSC 100-way vertebrate phastCons elements. ClinVar records were obtained from the January 2026 variant summary and VCV XML releases.

The ClinVar SNV and ClinVar indel benchmark datasets, including fixed splits and dataset cards, are shared in HuggingFace:

- Zygosity-aware ClinVar SNV benchmark: https://huggingface.co/datasets/scrc-dnai/clinvar-diploid-snv
- Zygosity-aware ClinVar indel benchmark: https://huggingface.co/datasets/scrc-dnai/clinvar-diploid-indel

The compound-heterozygous benchmark uses variants derived from HGMD Professional. The benchmark-generation code, split logic and evaluation implementation will be released, but HGMD-derived variant records will not be redistributed unless permitted by the applicable licence. A reconstruction procedure for appropriately licensed users is provided.

### 4.13 Code Availability

Diploid token definitions, preprocessing and local-alignment code, continual-pretraining scripts, the Contrastive Phase Loss implementation, GFMBench-API task implementations, fixed evaluation configurations and trained model checkpoints (NTv3-8M and NTv3-100M diploid and haploid variants across tokenizer and CPL ablation configurations) are available at https://github.com/scrcdnaimax/DNT-Diploid-Genomic-Foundation-Model. The release will identify the exact commit, software environment and checkpoint hashes used for each reported result.

## Acknowledgements

The authors thank the Kahn Family Foundation; the Flight Attendant Medical Research Institute (FAMRI); the Varda and Boaz Dotan Research Center in Hemato-Oncology, Tel Aviv University; the Ernest and Bonnie Beutler Research Program; and the Israel Innovation Authority, for their continuous support of our research.

## A Extended data

**Extended Data Fig. 1.**
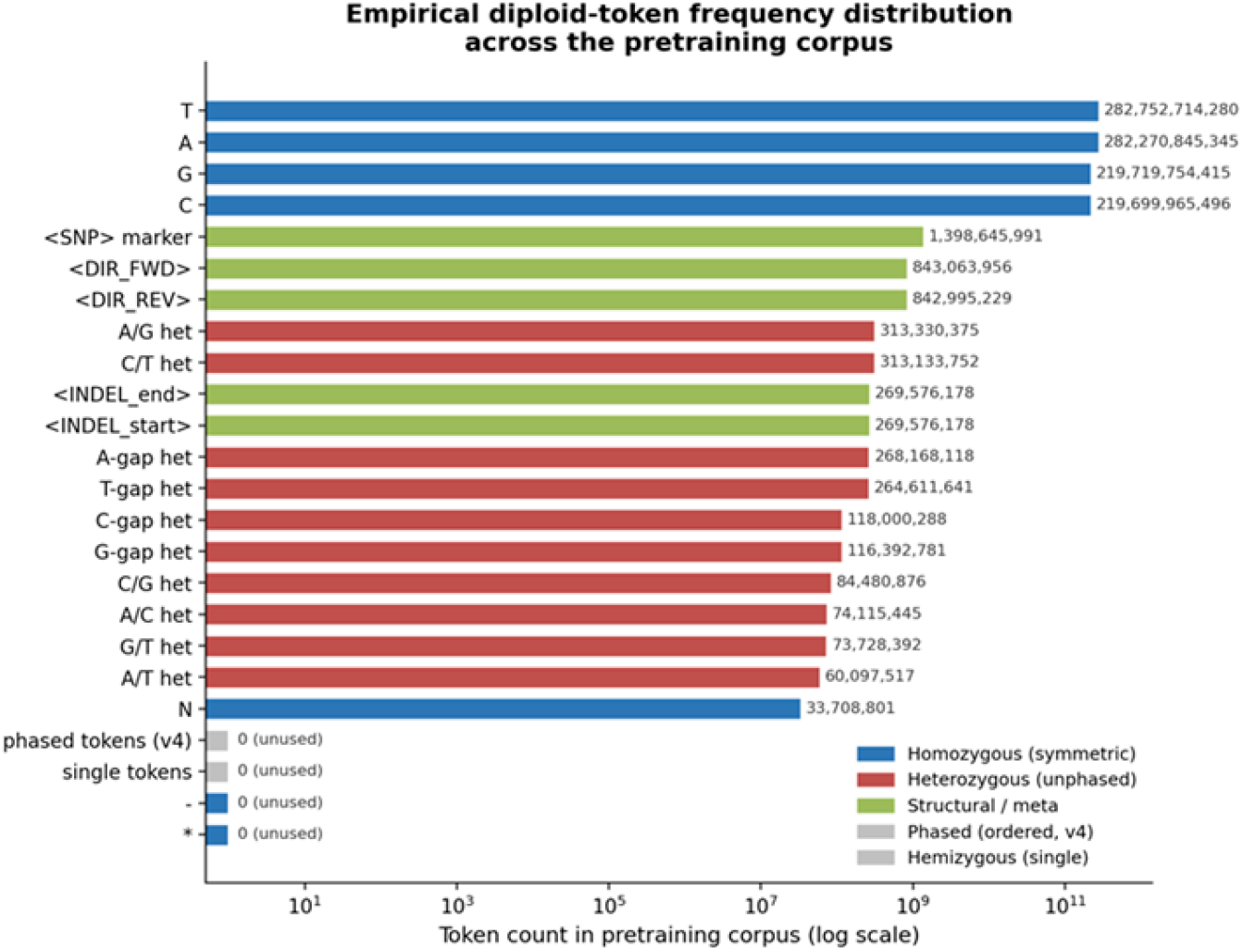
Empirical diploid-token frequency distribution across the pretraining corpus. Token counts across the pretraining corpus (log-scale x-axis). Homozygous reference-allele tokens (A/C/G/T) dominate by several orders of magnitude (2.2-2.8 × 10^11 each), as expected for reference-aligned diploid sequence. Among non-reference tokens, the <SNP> and <DIR_FWD>/<DIR_REV> structural markers and the ten unphased heterozygous symbols are all present at substantial frequency (10^8-10^9 tokens), whereas the phased/ordered and hemizygous token classes have zero observed counts. This confirms that the released pretraining corpus snapshot was tokenized with the unphased-symbol-plus-directional-marker scheme, with the phased and hemizygous vocabularies defined but not exercised in this corpus build.

**Extended Data Fig. 2.**
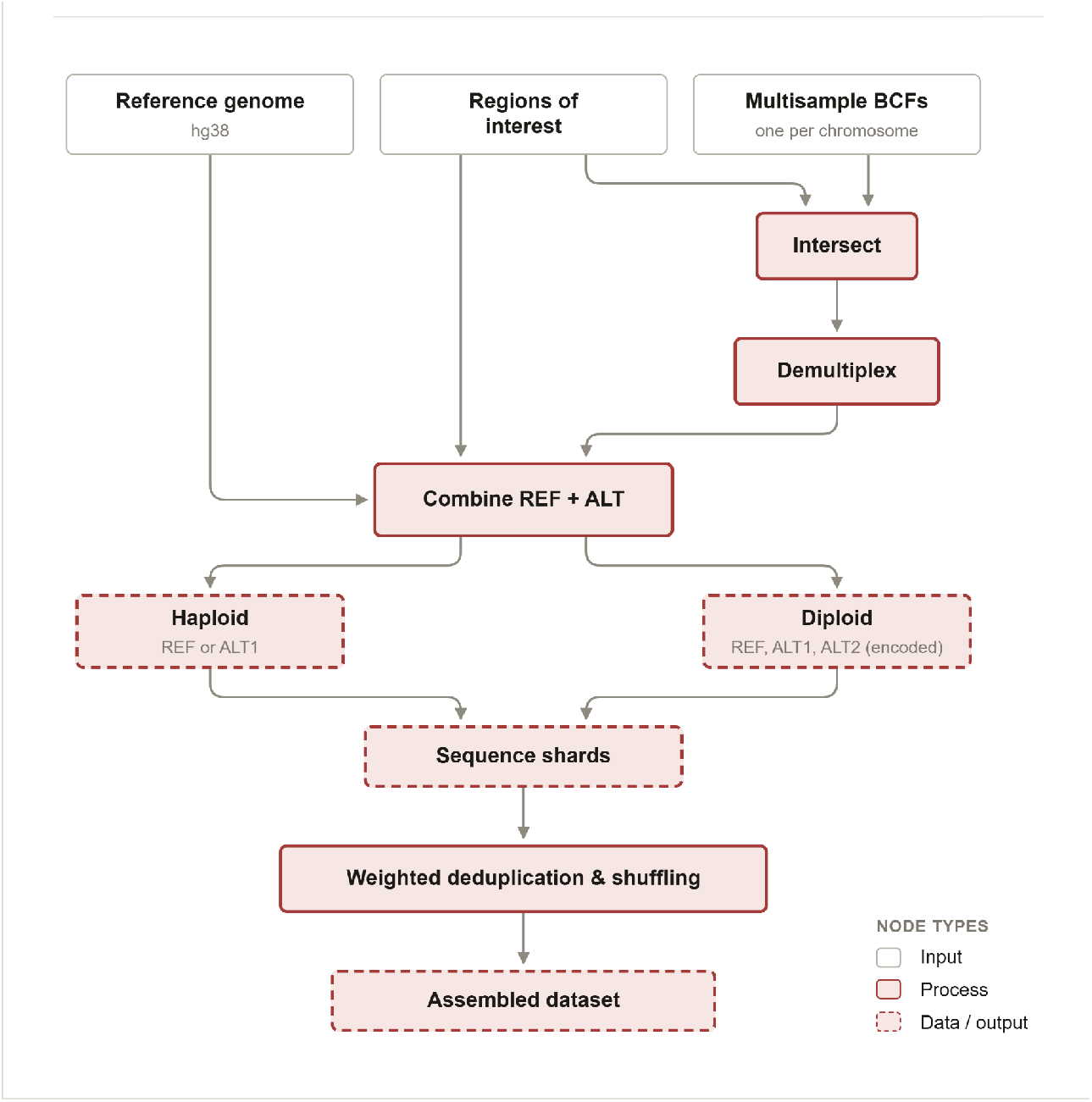
From raw phased VCFs to training-ready diploid tokens. Multisample BCFs, regions of interest, and the reference genome are intersected, demultiplexed, and combined into haploid and diploid REF/ALT sequences, then deduplicated and shuffled into the assembled dataset.

**Extended Data Table 1.** Pretraining interval sources and filtering criteria.

| Category | Criterion | Setting | Description |
| --- | --- | --- | --- |
| Interval source | GENCODE v47 Basic coding exons | - | Protein-coding exon (CDS) intervals drawn from the GENCODE v47 *Basic* transcript set, which retains representative, full-length transcript models. Supplies the coding-sequence component of the interval pool. |
| Interval source | phastConsElements100way | Log-odds score > 500 | Conserved elements from the UCSC 100-way vertebrate phastCons alignment. Only elements scoring above 500 are retained, selecting the most strongly conserved regions as the conservation component. |
| Filter | Minimum element length | $\geq 10$ bp | Elements and exons shorter than 10 bp are discarded to remove fragmentary or noise intervals. |
| Processing | Context-window inflation | 4,094 bp | Each retained interval is padded with a flanking genomic sequence out to a 4,094 bp window, giving the model surrounding sequence context around every core region. |
| Processing | Merge distance | 300 bp | Inflated intervals that overlap or lie within 300 bp of one another are merged into single contiguous intervals, reducing redundancy and fragmentation. |
| Filter | Canonical-chromosome filtering | chr1-22, X, Y | Intervals are restricted to the canonical assembled chromosomes; unplaced/unlocalized scaffolds and alternate/patch contigs are excluded. |
| Outcome | Merged coverage | 943 Mb (~30% of hg38) | Total span of the final merged interval set - 943 Mb, roughly 30% of the ~3.1 Gb hg38 genome. |

Sources, filters and processing steps defining the 140,099 continual-pretraining intervals spanning approximately 943 Mb (~30% of GRCh38).

**Extended Data Table 2.** HGMD seven-scenario zygosity and phase panel.

| HGMD scenario | MLM AUROC | +CPL AUROC | Delta (CPL – MLM) |
| --- | --- | --- | --- |
| Homozygous (zygosity) | 0.826 | 0.976 | +0.150 |
| Hemizygous (zygosity) | 0.813 | 0.992 | +0.179 |
| Carriers (zygosity) | 0.69-0.81 | 0.76-0.83 | +0.02 to +0.07 |
| Cis (phase) | 0.569 | 0.644 | +0.075 |
| Trans (phase) | 0.372 | 0.476 | +0.104 |
| Heterozygous (dominance) | 0.427 | 0.347 | -0.080 |

HGMD seven-scenario benchmark: AUROC for MLM and +CPL models across zygosity-driven (homozygous, hemizygous, carrier) and phase-driven (cis, trans) scenarios, plus a heterozygous dominance scenario as a phase/zygosity-independent comparison point

**Extended Data Table 3.**
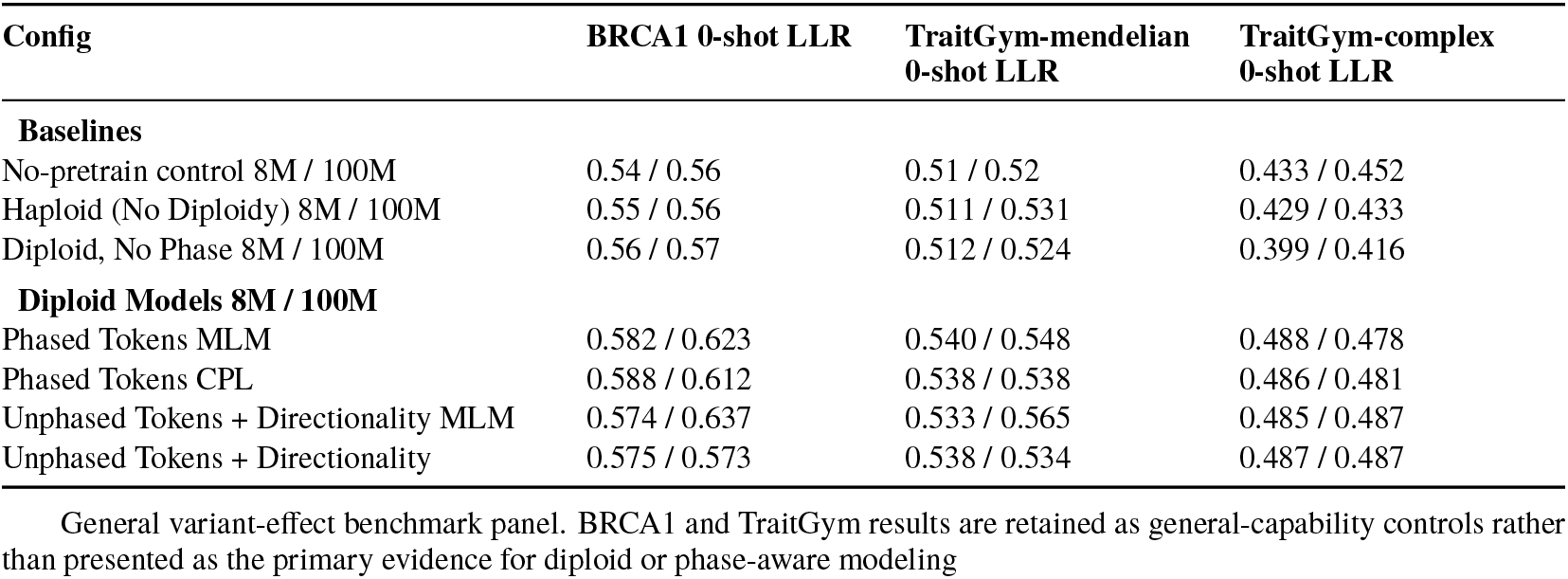
BRCA1 and TraitGym zero-shot general-capability controls.

**Extended Data Table 4.** Vocabulary size and pretraining-corpus token counts for the diploid tokenizers, with and without directionality markers, and for the haploid control.

|  | **Diploid with directional** | **Diploid w/o directional** | **Haploid** |
| --- | --- | --- | --- |
| Distinct symbols | 20 | 18 | 8 |
| Base tokens | 1,328,556,051,985 | 1,152,020,755,379 | 1,000,077,967,094 |
| Marker tokens | 4,765,026,739 | 2,224,090,029 | 1,735,823,006 |
| Non-ACGT base-token share | 0.17% | 0.15% | 0% |

## B Supplementary

### B.1 Supplementary Note 1: Extended comparison with prior genomic language models

Sequence pretraining. Early BERT-style DNA models such as DNABERT[5] introduced masked-language pretraining over k-merized genomic sequence, followed by more efficient tokenization and multi-species extensions in DNABERT-2[6]. Nucleotide Transformer scaled transformer pretraining across human and multispecies genomes and demonstrated broad downstream transfer[7]. These models established DNA language modeling as a useful pretraining paradigm, but they operate on haploid or allele-substituted sequences rather than native diploid genotypes.

Long context and alternative training paradigms. HyenaDNA[8] uses implicit long convolutions to model up to million-token genomic contexts at single-nucleotide resolution. Mamba[9] introduced selective state-space models with linear-time sequence scaling, and Caduceus[23] adapted this family to bidirectional, reverse-complement-equivariant DNA modeling. Evo[11] and Evo 2[12] further expanded long-context biological sequence modeling using Hyena-derived architectures. Genos[34] is a human-centric mixture-of-experts genomic foundation model optimized for 1 Mb sequence context. JEPA-DNA introduces a model-agnostic latent predictive training framework that complements token-level generative objectives, although its reported sequence inputs remain haploid[35]. These architectures address context length and reverse-complement symmetry, but they remain orthogonal to the problem addressed here: representing the two homologous alleles of an individual genome within a single sequence-native input. Caduceus in particular processes paired sequence views, but the pair reflects the two complementary DNA strands, not the two inherited homologues.

Variant- and genotype-centric models. SNP2Vec and related variant-centric models learn representations over SNP sequences or genotype-derived tokens rather than reference DNA alone[17, 18]. They include both alleles directly and support SNVs, but they use a single token to represent any indel, at any length, and therefore lose the actual mutation content. MutBERT[21] and UKBioBERT[22] model population-level or individual-level variant information to improve functional-genomics representations; MutBERT retains the underlying sequence and represents each position as a nucleotide probability distribution, thereby encoding where the genome is polymorphic rather than which two alleles a specific individual carries, so it does not directly model individual zygosity or phase. Genotype-imputation models such as STICI[19] and GenoBERT[20] show that transformer-style architectures can learn linkage disequilibrium and haplotype structure from genotype data; STICI separates the maternal and paternal copies before imputation, so zygosity and phase are captured directly. However, because these models act on discrete variant tokens over fixed marker panels and are optimized primarily for imputation, they do not retain the surrounding nucleotide sequence, do not target clinical variant-effect prediction, and generally support only predefined variant alternatives.

Sequence-to-function and evolutionary models. Enformer[10] and Borzoi[16] learn to predict regulatory and transcriptomic outputs from long DNA sequences and achieve strong noncoding varianteffect prediction; AlphaGenome[15] extends this paradigm with longer context and multimodal genomic tracks. These models evaluate variant effects by substituting alleles into a single sequence and comparing predictions, which is powerful for regulatory prediction but does not natively encode the realized diploid genotype of an individual. GPN-MSA[13] incorporates multiple-sequence alignments across vertebrate species to capture conservation and substantially improves genome-wide variant-effect prediction, and related work adds pretraining targets such as evolutionary conservation or functional tracks[14, 28]. These methods demonstrate the value of structured biological priors, but they describe cross-species conservation rather than within-individual diploid structure and are therefore orthogonal to modeling zygosity and inheritance within human genomes.

Zygosity-aware DNA language modeling. Rashidy et al.[25] and Saadat et al.[24] evaluate zygosity-aware DNA language modeling for genetic variation, ancestry and gene expression using dual inference over independent alleles. These works show that explicitly incorporating zygosity improves downstream prediction. However, treating the two alleles as separate sequence inputs requires two encoder passes and leaves joint interpretation of the allele pair to a downstream combination step, rather than representing both alleles together at each locus from the outset. Because checkpoints for the SNP2Vec-family models are not publicly released, they could not be benchmarked directly; we therefore adopt the dual-allele strategy of these works and refer to it as the ”biallelic” baseline mode.

Summary of the gap. Taken together, existing genomic language models show that sequence pretraining learns useful representations, that long-context architectures improve genomic modeling, that evolutionary priors add predictive signal, and that human variation carries phenotype-relevant information. Nucleotide sequence context and diploid genotype structure have each been modeled effectively, but not jointly within a single per-locus input representation. The remaining gap is native representation of the realized diploid genotype (zygosity, allele dosage, phase, and explicit indel sequence at each locus) in one sequence-native input. This work targets that gap.

**Supplementary Table 1.**
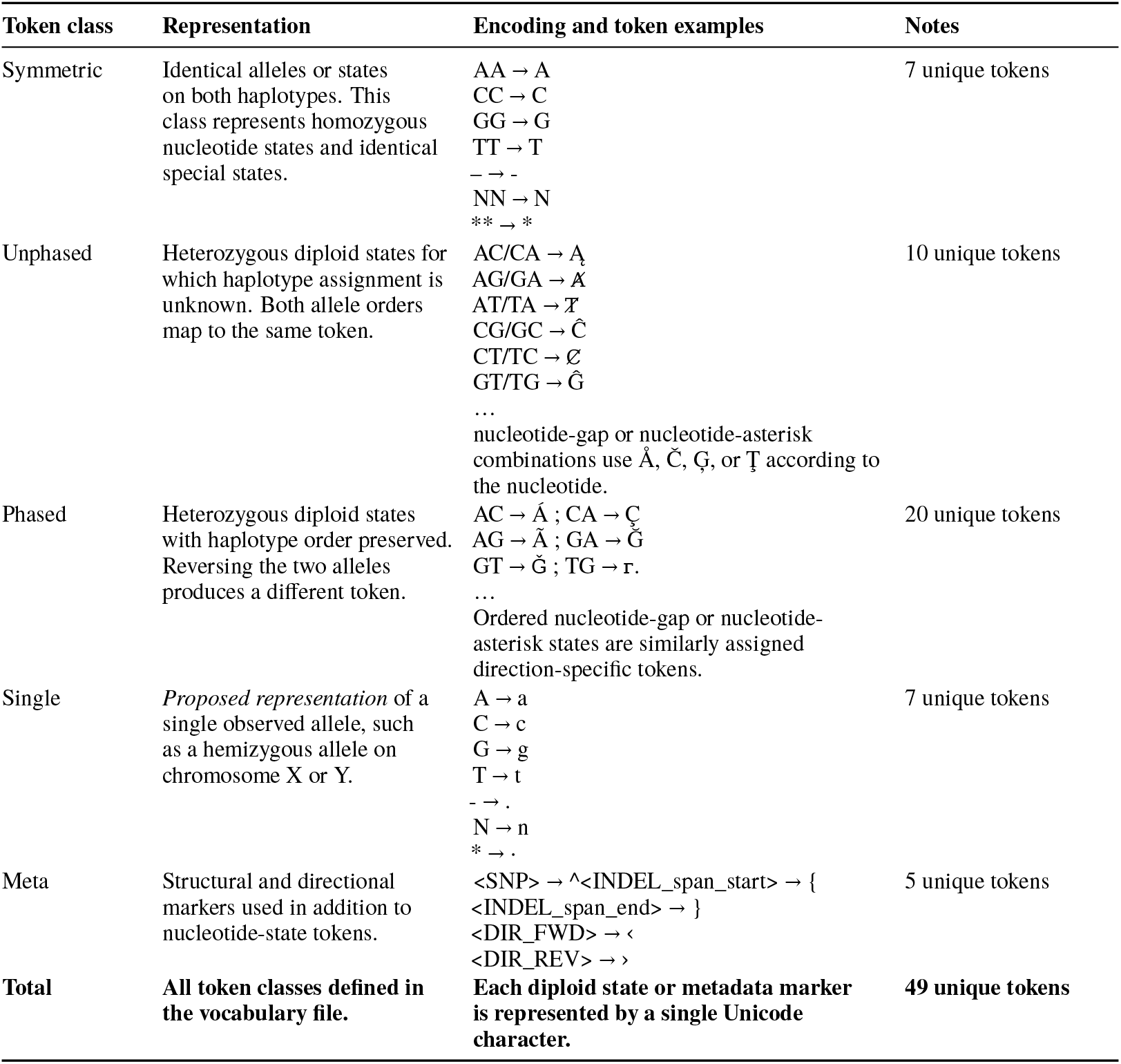
Diploid token vocabulary.

**Supplementary Table 2.**
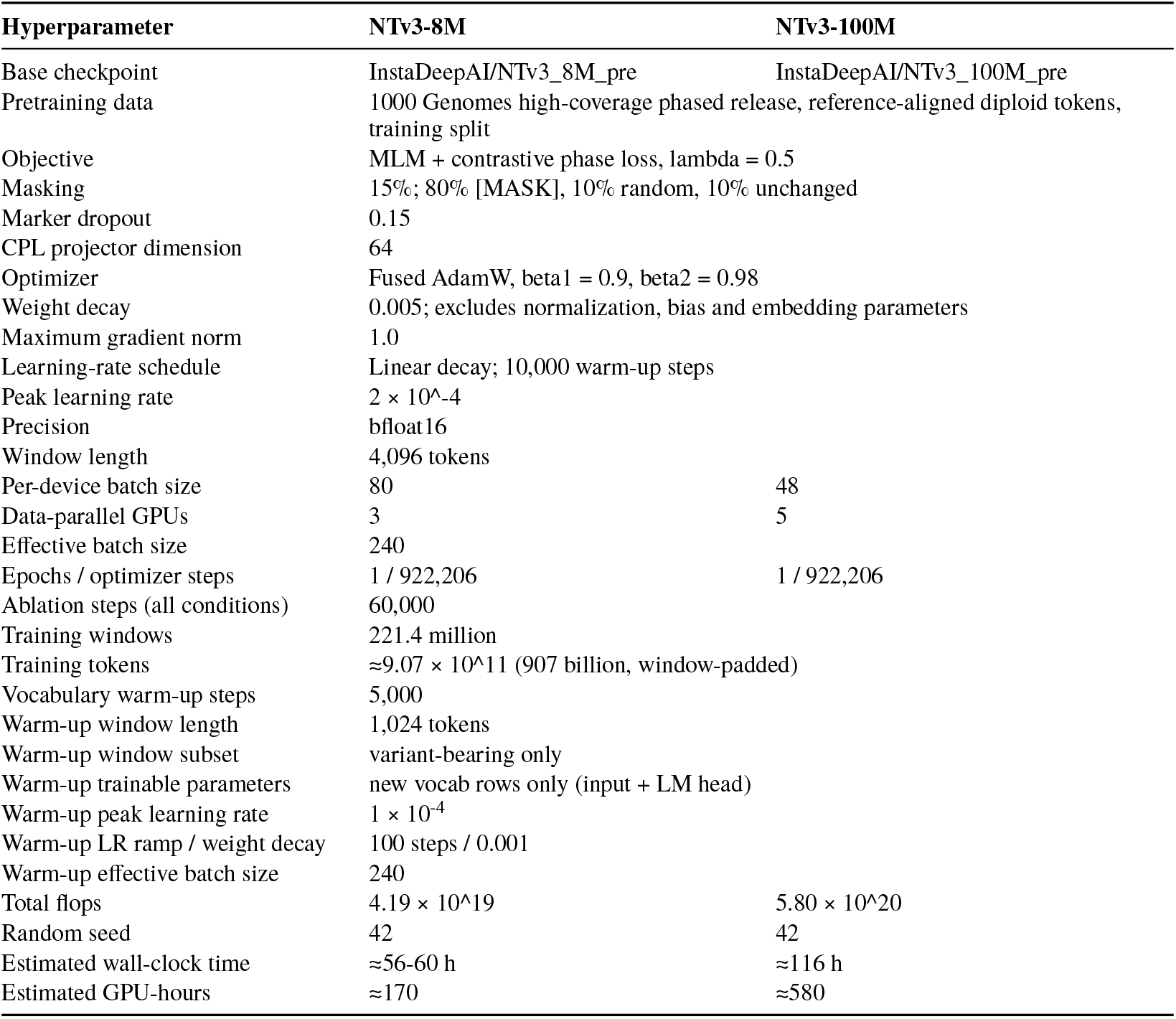
Continual pretraining configuration.

| Hyperparameter | NTv3-8M | NTv3-100M |
| --- | --- | --- |
| Base checkpoint | InstaDeepAI/NTv3_8M_pre | InstaDeepAI/NTv3_100M_pre |
| Pretraining data | 1000 Genomes high-coverage phased release, reference-aligned diploid tokens, training split |  |
| Objective | MLM + contrastive phase loss, lambda = 0.5 |  |
| Masking | 15%; 80% [MASK], 10% random, 10% unchanged |  |
| Marker dropout | 0.15 |  |
| CPL projector dimension | 64 |  |
| Optimizer | Fused AdamW, beta1 = 0.9, beta2 = 0.98 |  |
| Weight decay | 0.005; excludes normalization, bias and embedding parameters |  |
| Maximum gradient norm | 1.0 |  |
| Learning-rate schedule | Linear decay; 10,000 warm-up steps |  |
| Peak learning rate | $2 \times 10^{-4}$ | |
| Precision | bfloat16 |  |
| Window length | 4,096 tokens |  |
| Per-device batch size | 80 | 48 |
| Data-parallel GPUs | 3 | 5 |
| Effective batch size | 240 |  |
| Epochs / optimizer steps | 1 / 922,206 | 1 / 922,206 |
| Ablation steps (all conditions) | 60,000 |  |
| Training windows | 221.4 million |  |
| Training tokens | $\approx 9.07 \times 10^{11}$ (907 billion, window-padded) | |
| Vocabulary warm-up steps | 5,000 |  |
| Warm-up window length | 1,024 tokens |  |
| Warm-up window subset | variant-bearing only |  |
| Warm-up trainable parameters | new vocab rows only (input + LM head) |  |
| Warm-up peak learning rate | $1 \times 10^{-4}$ | |
| Warm-up LR ramp / weight decay | 100 steps / 0.001 |  |
| Warm-up effective batch size | 240 |  |
| Total flops | $4.19 \times 10^{19}$ | $5.80 \times 10^{20}$ |
| Random seed | 42 | 42 |
| Estimated wall-clock time | $\approx 56$-60 h | $\approx 116$ h |
| Estimated GPU-hours | $\approx 170$ | $\approx 580$ |

**Supplementary Table 3.** Training objective, budget and learning rate.

| Configuration | Frozen 8M | Frozen 100M | Fine-tuned 8M | Fine-tuned 100M |
| --- | --- | --- | --- | --- |
| *Training objective, 60,000 steps* |  |  |  |  |
| Marker-phase, MLM | $0.548 \pm 0.005$ | $0.540 \pm 0.020$ | $0.566 \pm 0.023$ | $0.560 \pm 0.014$ |
| Marker-phase, MLM + CPL | $0.555 \pm 0.016$ | $0.621 \pm 0.005$ | $0.610 \pm 0.008$ | $0.630 \pm 0.014$ |
| *Training budget, MLM + CPL at $2 \times 10^{-4}$* |  |  |  |  |
| 60,000 steps (166 genome-equivalents) | $0.555 \pm 0.016$ | $0.621 \pm 0.005$ | $0.610 \pm 0.008$ | $0.630 \pm 0.014$ |
| **922,206 steps (full epoch)** | **$0.613 \pm 0.009$** | **$0.624 \pm 0.005$** | **$0.649 \pm 0.005$** | **$0.639 \pm 0.005$** |
| *Learning rate, MLM + CPL at full epoch* |  |  |  |  |
| $5 \times 10^{-5}$ | $0.551 \pm 0.009$ | $0.610 \pm 0.011$ | $0.627 \pm 0.010$ | $0.625 \pm 0.006$ |
| **$2 \times 10^{-4}$** | **$0.613 \pm 0.009$** | **$0.624 \pm 0.005$** | **$0.649 \pm 0.005$** | **$0.639 \pm 0.005$** |

Compound-heterozygous *cis* versus *trans* classification; test AUROC on the locked 1,811-example split, mean ± s.d. over three probe seeds. Chance = 0.5. Rows within each block differ in one factor only. The learning-rate block is the largest single effect in this table: at 8 million parameters, training at $5 \times 10^{-5}$ costs 0.062 frozen AUROC relative to $2 \times 10^{-4}$ at the same budget, so schedule rather than objective is the dominant factor at full budget. The full-epoch rows are not budget-matched to the 60,000-step rows and the comparison between blocks is confounded by total compute.

**Supplementary Table 4.**
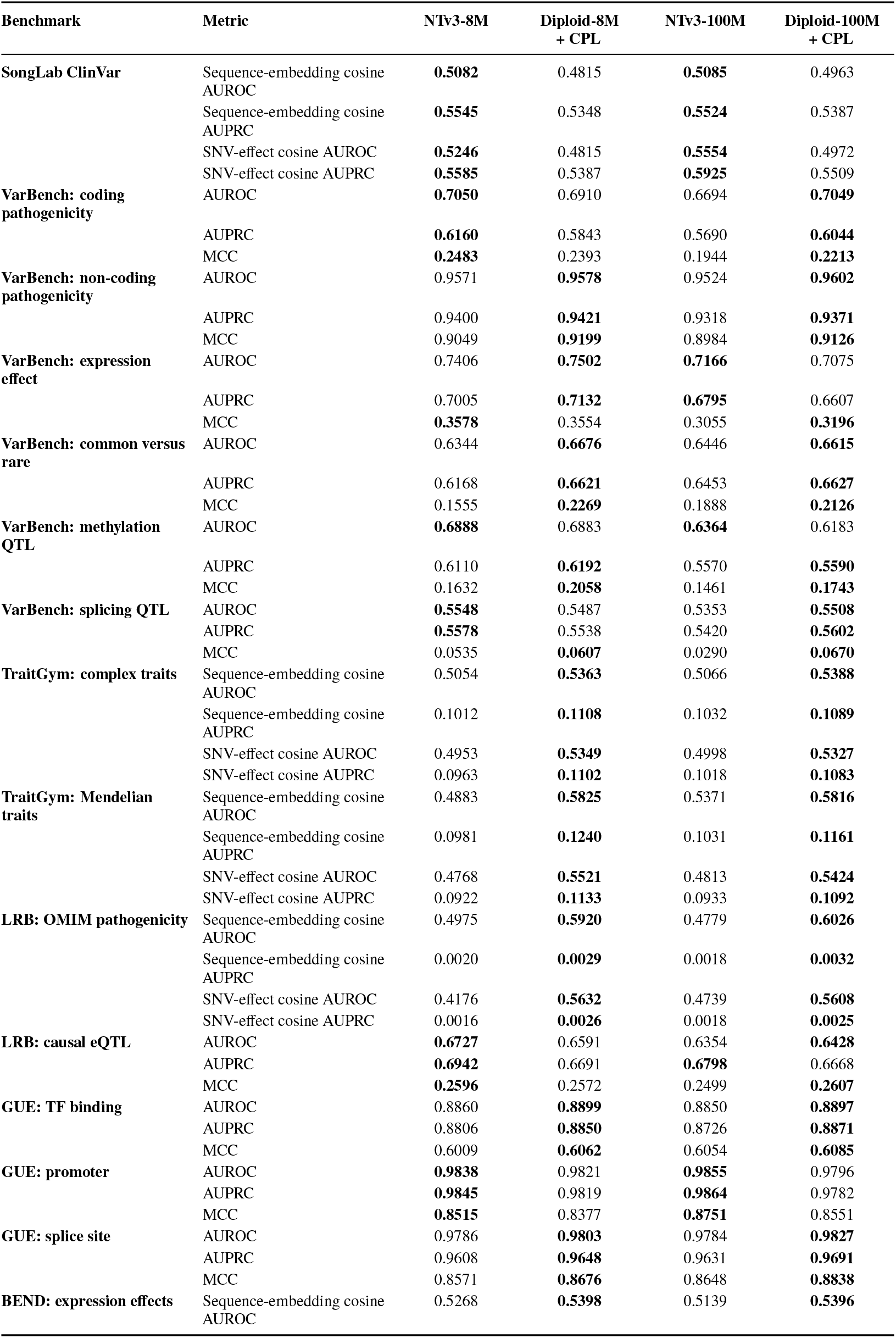
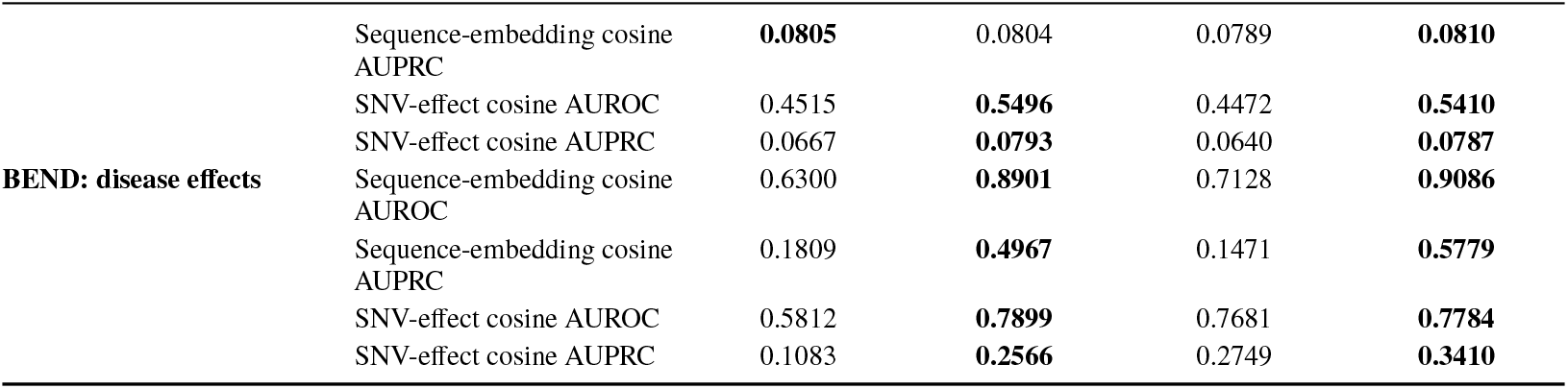
GFMBench-API general capability controls The original NTv3-8M and NTv3-100M checkpoints are compared with the corresponding 8-millionand 100-millionparameter diploid checkpoints trained with the marker-phase tokenizer (unphased allele-pair tokens with explicit directionality markers) and Contrastive Phase Loss (CPL). For supervised tasks, AUROC, AUPRC and Matthews correlation coefficient (MCC) are reported. For zero-shot tasks, sequence-embedding cosine and SNV-effect cosine AUROC and AUPRC are reported.Higher values are better; within each model scale, the higher value is shown in bold. Values are rounded to four decimals. Across the 54 selected benchmark-metric combinations, the diploid checkpoint exceeded its same-scale NTv3 baseline in 36 comparisons at 8M and 43 comparisons at 100M.

**Supplementary Table 5.** Contrastive Phase Loss weight.

| Run | $\lambda$ (constant) | Initialised from | Steps | Compound-het AUROC |
| --- | --- | --- | --- | --- |
| *Single-stage: the budget-matched $\lambda$ sweep* |  |  |  |  |
| MLM only | 0.00 | vocabulary-adapted baseline | 40,000 | 0.5473 |
| MLM + CPL, half weight | 0.25 | vocabulary-adapted baseline | 40,000 | 0.5963 |
| MLM + CPL | 0.50 | vocabulary-adapted baseline | 40,000 | 0.6079 |
| MLM + CPL, double weight | 1.00 | vocabulary-adapted baseline | 40,000 | 0.6162 |
| *Two-stage: not budget-matched to the rows above* |  |  |  |  |
| CPL withdrawn | 0.00 | MLM + CPL, genome-100 | 18,000 | 0.5963 |
| CPL introduced | 0.50 | MLM only, genome-100 | 18,000 | 0.6002 |

8M marker-phase models; effective batch 384, learning rate $2 \times 10^{-4}$ , seed 42. Compound-heterozygous full fine-tune, test AUROC, single run per cell. $\lambda$ , step budget and initialisation were read from each run's stored configuration. $\lambda$ is constant within every run; none of these are annealing schedules. The single-stage sweep is monotone in $\lambda$ ( $0.547 \rightarrow 0.596 \rightarrow 0.608 \rightarrow 0.616$ ), so $\lambda = 1.0$ is the highest cell rather than the $\lambda = 0.5$ carried into the main models; with one run per cell the 0.608 versus 0.616 difference lies inside the run-to-run spread observed elsewhere, and $\lambda = 0.5$ was fixed before this sweep was run. The two-stage rows add 18,000 steps on top of a stage-one checkpoint and are therefore not comparable to the single-stage rows.

**Supplementary Table 6.**
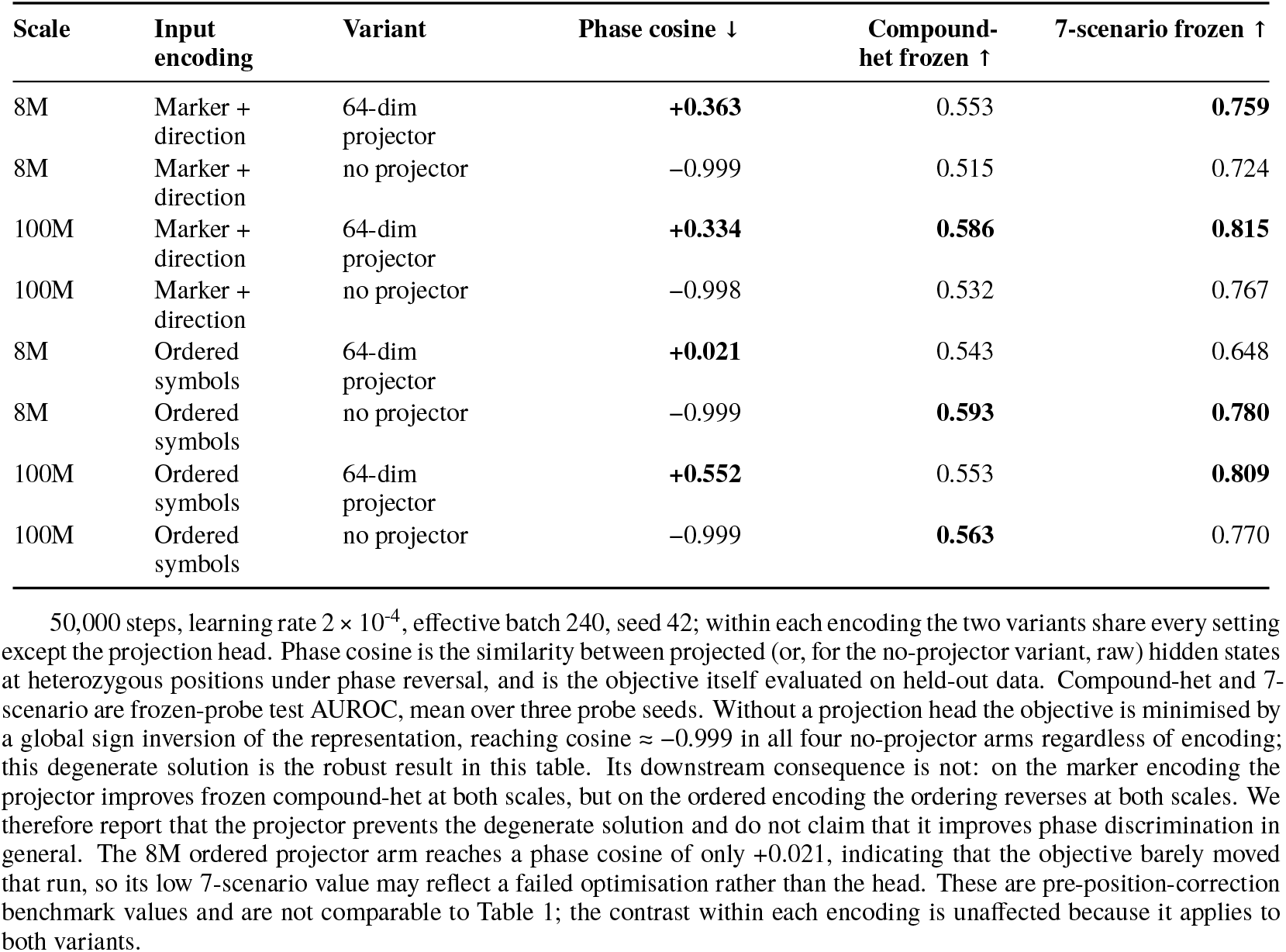
Projection head ablation for the Contrastive Phase Loss.

**Supplementary Table 7.** Input encoding, ordered symbols versus marker plus direction.

| Input encoding | Objective | Cis-trans frozen 8M | frozen 100M | fine-tuned 8M | fine-tuned 100M | 7-scenario 8M | 7-scenario 100M |
| --- | --- | --- | --- | --- | --- | --- | --- |
| Ordered symbols | MLM | 0.499 ± 0.010 | 0.529 ± 0.003 | 0.555 ± 0.007* | 0.534 ± 0.013* | 0.711* | 0.801* |
| Marker + direction | MLM | 0.501 ± 0.013 | 0.541 ± 0.003 | 0.528 ± 0.016 | 0.558 ± 0.023 | 0.712 ± 0.012 | 0.831 ± 0.003 |
| Ordered symbols | MLM + CPL | 0.543 ± 0.011 | 0.553 ± 0.004 | 0.592 ± 0.005* | 0.556 ± 0.001* | 0.648* | 0.809* |
| **Marker + direction** | **MLM + CPL** | **0.553 ± 0.015** | **0.586 ± 0.008** | **0.584 ± 0.007** | **0.592 ± 0.018** | **0.759 ± 0.005** | **0.815 ± 0.002** |

Test AUROC. The two encodings differ in how a heterozygous position is written: the ordered encoding assigns one atomic symbol per ordered allele pair, so the symbol itself carries phase, whereas the marker encoding uses an order-agnostic symbol plus a separate direction token. Both conditions are matched on optimizer steps (50,000), learning rate, effective batch, checkpoint lineage and evaluation harness, and each arm is evaluated under its own encoding, so the contrast isolates the encoding. marks cells whose runs do not carry distinct recorded seeds and are therefore run-to-run spread rather than seed replication.

**Supplementary Table 8.**
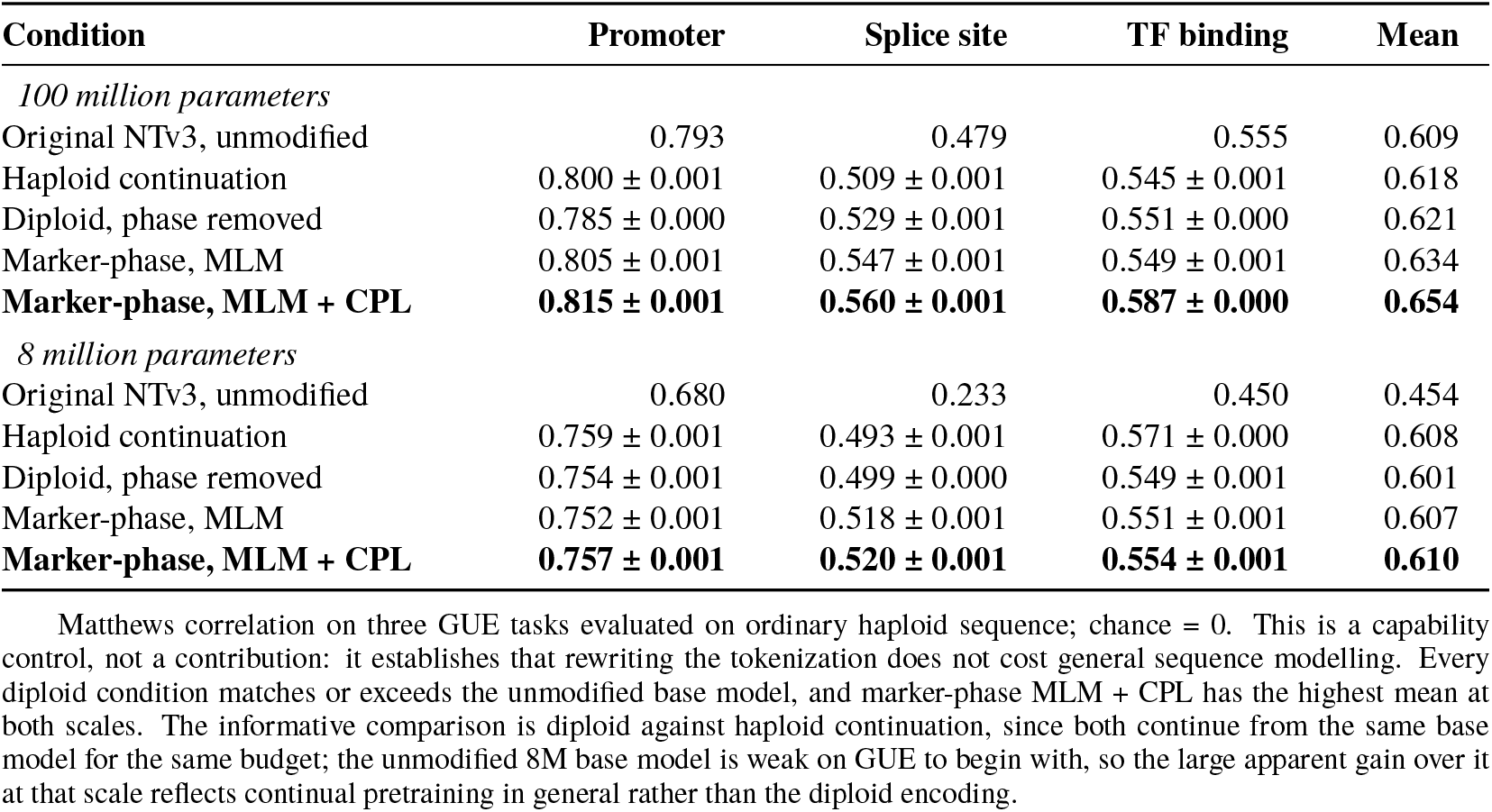
GUE sequence-modelling control.

## Footnotes

^1^https://hgdownload.cse.ucsc.edu/goldenpath/hg38/bigZips/hg38.2bit

